# Germline modifiers of the tumor immune microenvironment implicate drivers of cancer risk and immunotherapy response

**DOI:** 10.1101/2021.04.14.436660

**Authors:** Meghana Pagadala, Victoria H. Wu, Eva Pérez-Guijarro, Hyo Kim, Andrea Castro, James Talwar, Timothy Sears, Cristian Gonzalez-Colin, Steven Cao, Benjamin J. Schmiedel, Shervin Goudarzi, Divya Kirani, Rany M. Salem, Gerald P. Morris, Olivier Harismendy, Sandip Pravin Patel, Jill P. Mesirov, Maurizio Zanetti, Chi-Ping Day, Chun Chieh Fan, Wesley K. Thompson, Glenn Merlino, J. Silvio Gutkind, Pandurangan Vijayanand, Hannah Carter

## Abstract

With the continued promise of immunotherapy as an avenue for treating cancer, understanding how host genetics contributes to the tumor immune microenvironment (TIME) is essential to tailoring cancer screening and treatment strategies. Approaches that intersect SNP modifiers of molecular phenotype, such as gene expression, with disease phenotypes have shown promise for implicating causal genetic factors. Here we evaluated 194 literature-curated TIME associations and 890 associations detected with 157 immune phenotype (IP) components found using genotypes from over 8,000 individuals in The Cancer Genome Atlas. Of these 1084, 233 associations comprising 219 unique TIME-SNPs were also cancer relevant, associating with cancer risk, survival, and/or immunotherapy treatment response. Many cancer relevant TIME-SNPS overlapped regions of active transcription, and were associated with gene expression in specific immune cell subsets, such as macrophages and dendritic cells. TIME-SNPs associated with cancer risk and response to immunotherapy implicated genes involved in antigen presentation, especially by antigen presenting cells. The strongest associations with survival were with *PD-L1* and *CTLA-4*, suggesting that SNPs modifying the potential for immune evasion could contribute to disease progression. To assess whether our approach could reveal novel cancer immunotherapy targets, we inhibited *CTSS,* a gene implicated by cancer risk and immunotherapy response-associated TIME-SNPs; CTSS inhibition resulted in slowed tumor growth and extended survival *in vivo*. These results validate the potential of cancer relevant TIME-SNPs to implicate target genes for countering immune suppressive characteristics of the TIME and set the stage for future host genetics analysis integrating germline variation and TIME characteristics.

**Significance:** A systematic screen for common germline variants associated with the tumor immune microenvironment across > 8000 tumors reveals novel cancer risk factors and targets for immunotherapy.

## Introduction

Cancer is a disease characterized by heterogeneous somatic and germline mutations that promote abnormal cellular growth, evasion from the immune system, dysregulation of cellular energetics, and inflammation^1–4^. Both inflammation and immune surveillance contribute to the selective forces that shape tumor evolution^3–6^. Immunotherapies alleviating immune suppressive signals have emerged as a promising treatment strategy; however, response rates are low and the determinants of response remain elusive^7, 8^. Furthermore, the potential of galvanizing the immune system is still unmet due to an incomplete understanding of the complex tumor immune microenvironment (TIME). In particular, knowledge of germline factors and other intrinsic factors that interact with characteristics of tumors to render them sensitive to host-immunity or immunotherapy is lacking. Germline variation is responsible for a considerable proportion of variation in immune traits in healthy populations^9, 10^. In the context of tumors, germline variants are associated with immune infiltration, antigen presentation and immunotherapy responses^11, 12^. Autoimmune germline variants modify ICB response and variants underlying leukocyte genes predict tumor recurrence in breast cancer patients^13, 14^. For example, the common single nucleotide polymorphism (SNP) rs351855 in *FGFR4* was found to suppress cytotoxic CD8+ T cell infiltration and promote higher immunosuppressive regulatory T cell levels via increased *STAT3* signaling in murine models of breast and lung cancer^15^. Normal genetic variation underlying major histocompatibility complex molecules, MHC-I and MHC-II, dictate which mutations in an individual’s tumor can elicit immune responses, and play a role in antigen-driven host anti-tumor immune activity that influences tumor genome evolution through immune selection^16, 17^. Polymorphic variation in these regions has also been linked to treatment outcomes^18–20^. Recent literature highlights polymorphisms in other immune-related genes such as *CTLA-4*^21^*, IRF5*^22^ and *CCR5*^23, 24^ that also affect treatment outcomes.

Efforts to identify germline variation associated with anti-tumor immune responses have pointed to effects on immune infiltration levels and immune pathways, such as *TGF-β* and *IFN-ɣ*^11, 12, 25^. Genes with significant cis-eQTLs in the TCGA are both enriched for immune-related genes and associated with immune cell abundance within the TIME^26^. These studies provide evidence that variants may act through specific effects on immune cells. eQTL profiling of 15 sorted immune cell subsets from healthy individuals found that the effects of many eQTLs were specific to immune cell subsets^27^. Understanding mechanisms and cell-type effects of TIME host genetic interactions could not only identify aspects of immunity that negatively impact cancer and immunotherapy outcomes, but also point to putative targetable cell types and molecules for modulating immune responses.

Here, we sought to identify common germline variants associated with TIME characteristics that are also associated with cancer outcomes, reasoning that such dual associations would implicate the aspects of immunity most critical for tumor control and uncover putative new targets for immunotherapy^28, 29^. We combined a new analysis of The Cancer Genome Atlas (TCGA) with previous germline studies to identify 1084 TIME associations and further analyzed them to converge on a subset associated with cancer outcomes. Adaptive immune genes relating to antigen processing and presentation, including several class I and II MHC genes, were implicated in risk for multiple cancers in the UK Biobank and validated in independent cohorts. We also found immune checkpoint and Th17 variants underlying immune evasion to be associated with overall and progression-free survival, respectively. Lastly, we identified 11 SNPs associating with ICB response in publicly available cohorts. Since overlapping eQTL and GWAS signals can suggest causal associations, we selected one such gene, *CTSS,* for validation as a possible immunotherapy target and found that inhibition of CTSS prolonged survival and reduced tumor growth in an MC38 murine model. These results illuminate the role of common genetic variation underlying the TIME in cancer risk and survival while also providing a potential novel avenue for immunotherapy target discovery. The study design is summarized in **Figure 1A**.

**Figure 1:**
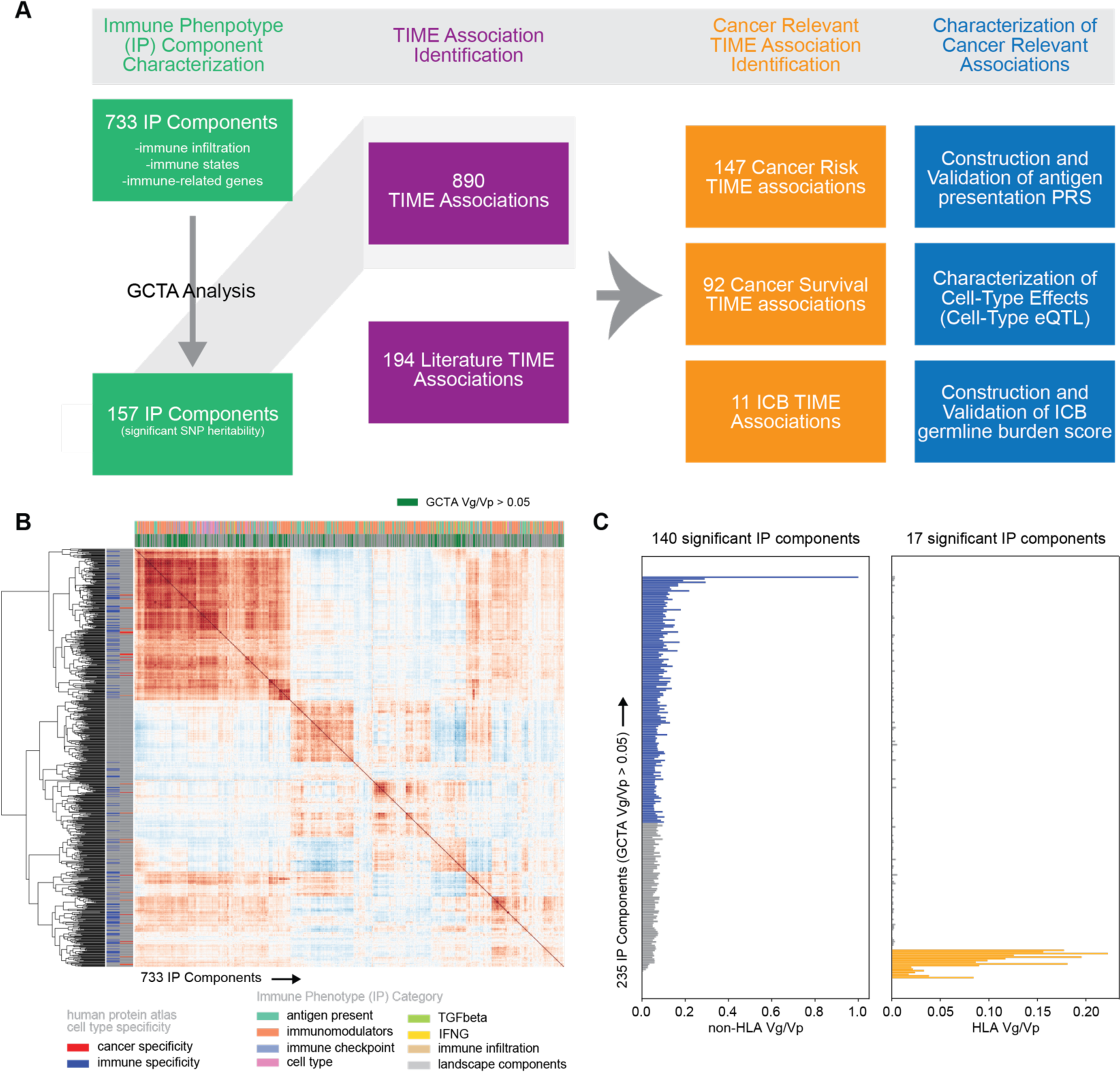
Characterization of Tumor Immune Microenvironment (TIME). **(A)** Overview of the TIME germline analysis. **(B)** Clustermap depicting 733 IP components and their pairwise correlation across 30 tumors in the TCGA. **(C)** Horizontal barplot of variance in phenotype explained by variance in genotype (Vg/Vp) for 235 IP components estimated separately genome-wide excluding the HLA locus (left panel) and using only the HLA locus (right panel).

### Identifying heritable characteristics of the tumor immune microenvironment (TIME)

To focus on common germline genetics with the potential to modify tumor immune responses, we assessed which characteristics of the TIME showed evidence of SNP heritability. To describe the TIME, we collected a comprehensive set of immune phenotype (“IP”) components comprising composite measures derived from bulk gene expression and expression levels of individual immune-related genes (**Figure 1B**). Composite phenotypes included infiltrating immune cell levels calculated using CIBERSORTx (immune infiltrates) and 6 immune subtype scores from a pan-cancer TCGA analysis by Thorsson et al. (landscape components). Immunomodulators were collected from Thorsson et al., where weighted gene correlation network analysis was used as an unbiased systematic approach to identify gene sets relevant to the TIME. We included genes from these sets along with immune checkpoint genes, cell type markers, antigen presentation genes, TGF-β pathway genes, and IFN-γ genes as these have been implicated as important modifiers of the TIME. After removing IP components with high numbers of zero values to reduce spurious associations, we retained 724 immune-related genes and 9 composite phenotypes (733 IP components total) measured across 30 cancer types **(Supplementary Table 1**, **Figure S1)**.

We evaluated the potential of germline variation to explain inter-tumor differences in IP components by performing SNP heritability analysis **(Figure 1A)**. Since highly polymorphic regions such as the HLA locus can inflate SNP heritability estimates, we separately estimated SNP heritability attributable to the HLA locus and the rest of the genome. We identified 235 (32.0%) IP components where levels were SNP-heritable (>5% of variance in expression or composite value was attributable to genetic variance; *i.e.* Vg/Vp > 5%; **Figure 1C; Supplementary Table 2**). For these 235 IP components, we conducted 2-state GCTA analysis and identified 140 (59.6%) that had a significant proportion of SNP heritability attributable to regions outside the HLA locus, while 17 (7.2%) were mostly attributable to the HLA locus at an FDR < 0.05. We focused our TIME-SNP discovery analysis on these 157 SNP-heritable IP components.

### Detecting putative germline modifiers of the tumor immune microenvironment

To build a comprehensive dataset of germline variants associated with the TIME, we integrated associations with SNP-heritable IP components and immune SNPs previously implicated by published studies. To identify genetic variants underlying IP component SNP heritability, we performed a genome-wide association study (GWAS). First, we performed GWAS for each of the 140 heritable IP components outside of the HLA locus across individuals of European ancestry in the TCGA **(Figure S2A)**. Only common germline variants with minor allele frequency > 1% were considered and imputation quality (Rsq) was evaluated to ensure high accuracy (**Figure S2B)**. No evidence of inflation was observed (**Figure S2C).** Using linkage and distance-based clumping^30^, we identified 825 associations with 75 SNP-heritable IP components at a threshold of 7.1x10^-8^ (Bonferroni-corrected suggestive threshold of 1x10^-5^), with 545 of the 825 (66.1%) passing a Bonferroni- corrected genome-wide significance threshold of 3.6x10^-10^ (**Figure 2A**, **Supplementary Table 3**). These 825 TIME associations implicated 795 unique TIME-SNPs. *Cis* associations, defined as an associated locus occurring within 1 MB of an IP component gene transcription start site, encompassed the majority (95.0%) of associations^31^, while 5.0% of the associations were *trans*. Mechanisms of *trans* associations are complex and tend to have weaker effects on transcriptional regulation^32^. In contrast, *cis* associations are proximal to an IP component and have more direct effects on transcription. Overall, *ERAP2* (181, 21.9%)*, CCBL2* (76, 9.2%)*, DHFR* (75 9.0%) and *ERAP1* (70, 8.5%) had the most germline associations **(Figure S2D)** of the 140 IP components tested.

**Figure 2:**
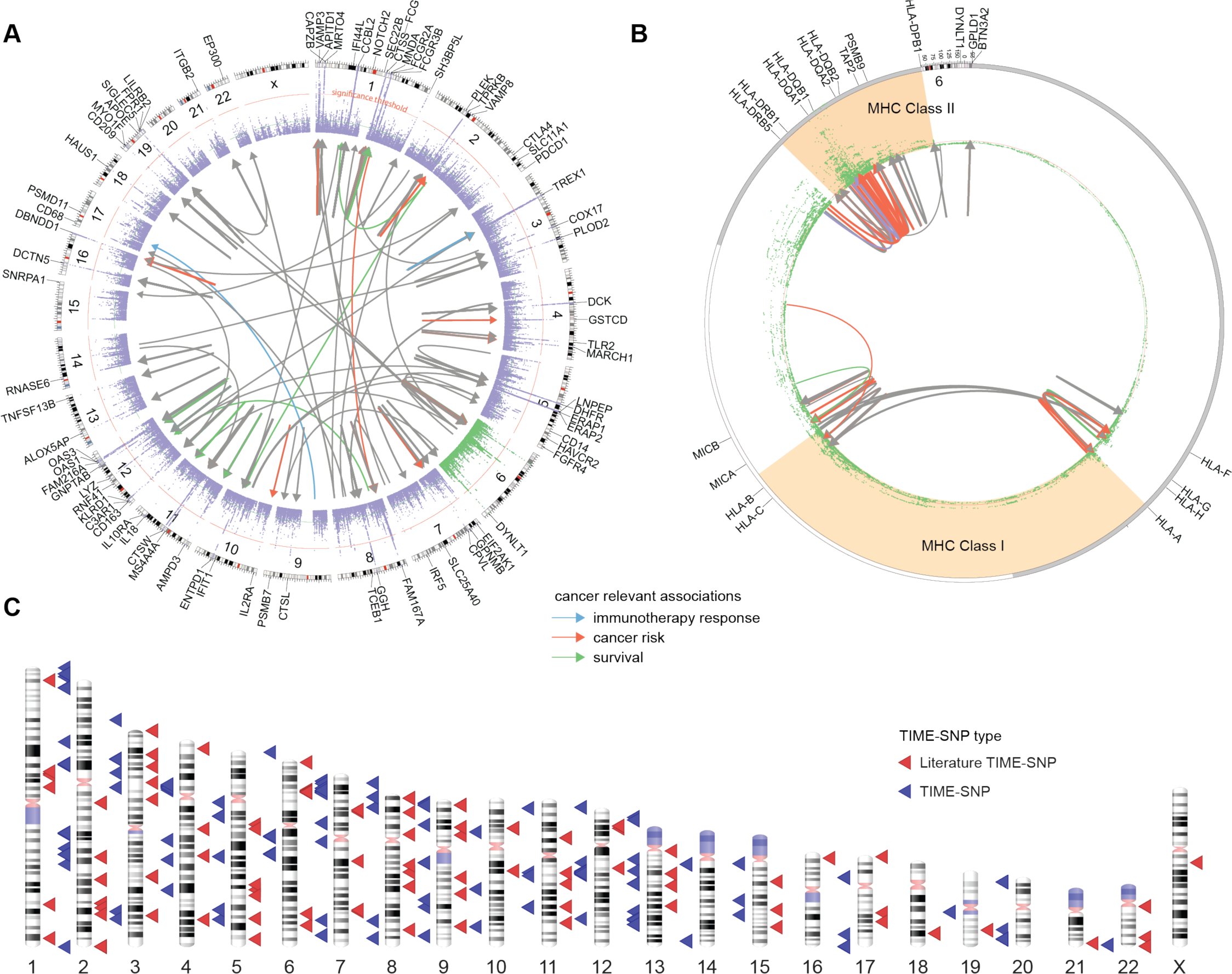
Detecting putative germline modifiers of the tumor immune microenvironment. **(A)** Locuszoom plot summarizing 890 associations between germline SNPs and 93 IP components. Outer ring represents locations of all 157 tested IP components. Links are colored if implicated in cancer risk (orange), survival (green) or immunotherapy response (blue) **(B)** Significant associations between SNPs and 17 IP components in the HLA region detected through conditional GWAS analysis for effects on gene expression using either a basic alignment to the reference genome (conditional), or allele specific expression obtained by aligning to a patient-specific HLA reference allele set. **(C)** Ideogram plot of TIME-SNPs implicated by our discovery analysis (red) and literature curation (blue).

To remove HLA region associations solely attributable to LD structure^33, 34^, we conducted conditional GWAS analysis for seventeen IP components corresponding to genes in the HLA region of chromosome 6, iteratively adding the strongest associated SNP on chromosome 6 as a covariate until no significant SNPs remained. Alignment to a general HLA gene reference can introduce error into expression level estimates due to the highly polymorphic nature of these genes. We therefore also revisited SNP associations with gene expression estimates derived from allele-specific RNA alignments^35^ **(Methods)** and performed GWAS analysis using allele specific expression. In total, 63 independent TIME-SNPs and 65 TIME associations were associated with HLA region gene expression **(Figure 2B)**; MHC Class II genes, *HLA-DQB1* and *HLA-DRB5,* had 8 and 7 significant LD-independent associations, respectively. Generally, LD-independent SNPs clustered by genomic regions with *HLA-A, HLA-B, HLA-C* associated variants falling in the MHC Class I genomic region and *HLA-DQB1, HLA- DQA1, HLA-DPB1, HLA-DRB5* associated variants falling in the MHC Class II genomic region **(Figure S2E)**. rs17612852 was associated with *HLA-DQB1* expression using both the reference alignment and allele-specific expression. Combining GWAS and conditional HLA GWAS associations, we identified 890 TIME associations and 858 unique TIME-SNPs.

We noted some correlation among IP components across tumors, especially for components associated with macrophages and lymphocytes which were among the most abundant infiltrating immune cells **(Figure S2F)**. We investigated whether IP component correlation would inflate the chance of detecting SNPs associated with a particular group, however analysis of summary statistics showed that despite their correlation, IP components typically did not recover the same SNP associations unless the IP components were genes encoded at the same genomic locus, such as *ERAP1* and *LNPEP* and *OAS1* and *OAS3* (**Figure S3F)**. Clustering of pearson correlation values amongst SNP-heritable IP components revealed 2 major groups of genes (**Figure S3A)**. The largest group included MHC Class I and II genes along with macrophage genes *VSIG4, CD163, FCGR2A FCGR3A, HAVCR2, LILRB2, LILRB4* and *CD53* **(Figure S3B)** and was most strongly associated with antigen presentation, dendritic cell processing, and *IL-10* production (**Figure S3C).** The next largest group comprised two anti-correlated subgroups of genes which contained *EP300* and *TREX1* respectively **(Figure S3D)**. This group of genes were related to innate immune activation, the C-type lectin receptor signaling pathway and antigen presentation **(Figure S3E)**. Two major groups of genes correlated with the top 2 principal components from Principal Component Analysis (PCA) conducted on the expression of the 157 unique SNP-heritable IP components across TCGA tumors. *CD53, CD86* and *CYBB*, which are highly correlated (ρ > 0.7) to the Thorsson et al.^36^ Macrophage Regulation score, were major contributors to PC1 while *HACD2, LNPEP* and *EP300,* were major contributors to PC2 **(Supplementary Table 4)**.

Previous studies of germline variation and important modulators of immune checkpoint response, such as *APOE*^37^*, CTSW*^38^*, CTLA-4*^21^*, PD-L1*^39, 40^*, PD-1*^41–43^*, CXCR3/CCR5*^23^*, IRF5*^22^ and *FGFR4*^15^ along with immune signatures and immune cell infiltration have been conducted^11, 26, 44^. We incorporated these 194 germline associations from literature into our analyses **(Figure 2C**, **Supplementary Table 5)**. Like Shahamatdar et al.^11^, we included immune infiltrates estimated from bulk RNA sequencing into the set of immune components we investigated, however, our filter on the proportion of phenotypic variance explained by genotype eliminated one of the two associations reported by Shahamatdar et al. which related to T follicular helper cell infiltration (V(g)/V(p) < 0.000001 whereas we required V(g)/V(p) >= 0.05). Zhang et al.^38^ took a fundamentally different approach, analyzing ER+ breast cancer-associated variants from Michailidou et al.^45^ for proximity to immunoinflammatory GWAS SNPs. The top SNP, rs3903072, was an eQTL for *CTSW* in breast cancer. Although not specifically focused on breast cancer, our study also identified *CTSW* as a SNP-heritable IP component (GCTA V(g)/V(p)= 12.1%) and detected a pan-cancer association with rs3903072 (beta = 0.21, p=2.8e-36). The study by Sayaman et al.^44^ focused on 139 immune traits described in the Thorsson et al.^36^ paper, of which 106 were immune signatures and 33 included immune measures such as TCR/BCR characteristics, CIBERSORTx infiltration and antigen load. Comparing gene results between Sayaman et al. and our study, 10 genes were shared between our analyses, *HLA-DRB5, HLA-B, HLA-DRB1, MICB, HLA-DQB1, HLA-DQB2, HLA-DQA1, HLA-DQA2, MICA, HLA-C*, emphasizing the importance of MHC Class I and II machinery in modifying the TIME. Of our variants, 31 were in linkage disequilibrium (LD) (R^2^ > 0.20) with 485 of 598 Sayaman et al. TIME-SNPs. Using R^2^ > 0.50, 19 of our variants were in LD with 361 Sayaman et al. TIME-SNPs.

Combining TIME-associations and literature SNPs resulted in a set of 1084 candidate SNPs. A number of TIME-SNPs were associated with multiple IP components; thus, we had a greater number of associations than TIME-SNPs. For example, within our own discovery pipeline, rs2693076 was associated with *LILRB2, PLEK, MYO1F,* and *CD14*. From literature curation, Sayaman et al. identified associations with rs2111485 and multiple signatures, including interferon-signaling and *IFIT3* signaling.

### Identification and characterization of TIME-SNPs related to cancer outcomes

A screen to detect TIME-SNPs would be expected to detect variants associated with immune traits more generally, including variants that may have little relevance to cancer. To focus our analysis on TIME-SNPs with evidence for cancer relevance, we evaluated their association with disease risk, progression and response to immunotherapy. SNPs associated with any of these aspects were considered cancer relevant.

SNPs were designated cancer risk associated if they had previously been implicated by a cancer GWAS study in the NHGRI-EBI GWAS catalog^46, 47^, Vanderbilt PheWAS catalog^48^ or were found to associate with an ICD10 code related to cancer in a PheWAS of the UK Biobank^49^ **(Methods)**. Rather than a GWAS which analyzes many genetic variants with one phenotype, a PheWAS analyzes many phenotypes compared to a single genetic variant. In our study, we conducted a PheWAS in the UK Biobank with TIME-SNPs and identified 61 cancer risk associations **(Supplementary Table 6)**, We also identified 64 cancer risk associations through intersection with the NHGRI-EBI GWAS catalog and 92 associations through intersection with the Vanderbilt PheWAS catalog. Although we used different methods to assess cancer risk, we observed high overlap in risk variants identified by the three sources **(Figure S4A)**. When assessing overlap based on the corresponding IP components, a higher degree of overlap was observed, with only 2 SNP-heritable IP components, *TAP2* and *LNPEP,* being uniquely implicated by the UK Biobank **(Figure S4B)**. In total, 138 unique TIME-SNPs associated with 41 IP components had cancer risk associations. This included risk associations with non-melanoma skin cancer, melanoma, lung cancer, prostate cancer, breast cancer and head and neck squamous cell carcinoma. Most cancer risk associations were found with the MHC II gene signature described in Sayaman et al.^50^ (31), *CTSS* expression (12) and *ERAP2* expression (11).

We next evaluated association of TIME-SNPs with overall and progression-free survival in the TCGA. We identified 92 associations and 87 variants that were significantly associated with overall or progression-free survival in at least one tumor type (FDR < 0.05) **(Supplementary Table 7)**. Of the variants associated with overall survival, the majority of associations occurred in cervical squamous cell carcinoma and endocervical adenocarcinoma (CESC, 8), uterine carcinosarcoma (UCS, 8) and thyroid cancer (THCA, 7). Variants associated with progression-free survival were also found in cervical squamous cell carcinoma and endocervical adenocarcinoma (CESC, 6), but otherwise were most frequent in stomach adenocarcinoma (STAD, 5), rectum adenocarcinoma (READ, 5), and liver hepatocellular carcinoma (LIHC, 5). Variants associated with *PD-1*, the MHC II signature, *GPLD1, ERAP1, ERAP2, CTSW*, dendritic cell signatures, *VAMP3* and *FAM216A* were associated with survival in more than 1 tumor type in the TCGA. For example, the intronic *PD-L1* variant, rs822339, was associated with overall survival in multiple cancer types, including lung adenocarcinoma (LUAD), kidney chromophobe cancer (KICH) and thyroid cancer (THCA), confirming survival associations reported by Yoshida *et al*^39^. Of these 87 variants, 5 remained significantly associated with survival in a Cox proportional-hazards analysis with covariates **(Methods)**, including 3 *PD-L1* variants, an *ERAP2* variant and a *DCTN5* variant **(Supplementary Table 7)**.

To investigate the implication of TIME-SNPs for immune checkpoint blockade (ICB) response, we collected sequencing and ICB response information for 276 patients with melanoma treated with immune checkpoint inhibitors from 4 studies^51–55^, and imputed SNPs from exome sequencing data. Accuracy of exome-based imputation was assessed by comparing original TCGA genotype calls to genotypes imputed in from TCGA exome data at positions matching those in the ICB data; aside from variants on chromosome 6 within the HLA region most variants were accurately imputed **(Supplementary Table 8).** Ultimately, of the 1084 TIME-SNPs, we considered 525 that could be imputed with sufficient quality (minor allele frequency > 0.05 in all 4 discovery ICB cohorts with imputation accuracy of at least 0.3^56, 57^. For ICB variant discovery, we conducted meta-analysis with METAL^58^ and found that 6 SNPs were significantly associated with ICB response (FDR < 0.25), implicating IP components such as *PSMD11, ERAP1, TREX1, ERAP2* and a T follicular helper cell signature. With a less stringent FDR threshold (FDR < 0.5), 11 variants were associated with ICB response, implicating the following additional genes: *CTSS, FAM216A, DHFR, DCTN5, LYZ* **(Supplementary Table 8)**.

In total, these analyses implicated 219 TIME-SNPs and 233 associations as cancer relevant. Assessing overlap based on SNPs passing significance thresholds, 12 variants associated with *DBNDD1, CCBL2, ERAP2, GPLD1, DCTN5, HLA-DQA1, HLA-DRB5, HLA-DQA1, CTLA-4* and the MHC II signature were implicated in both risk and survival analysis. The *CTSS* variant, rs2305814, was associated with cancer risk and ICB response, the *DHFR* variant, rs434130, was associated with cancer survival and ICB response and the *DCTN5* variant, rs546055 was associated with cancer risk, survival and ICB response. IP components *CTSS, ERAP1, TREX1, ERAP2* and *DCTN5* were implicated in both cancer risk and immunotherapy response while *FAM216A, ERAP1, TREX1, ERAP2, DHFR, DCTN5,* and *LYZ* were implicated in both cancer prognosis and immunotherapy response.

Focusing on these 219 cancer relevant TIME-SNPs, we sought to understand what aspects of the tumor- immune interface were affected. Cancer relevant TIME-SNPs were associated with 65 unique IP components. Several of these were antigen presentation and macrophage regulation genes, including both MHC I and II pathway genes **(Figure 3A)**. Fifty-two of the 233 (22.3%) associations were from literature curation and included associations with the MHC II signature, the T follicular helper cell signature, IFN, the IFIT3 attractor signature, *CTLA-4*, *PD-1*, *PD-L1*, *APOE*, *CTSW*, and monocyte, dendritic cell and TH2 cell infiltration. Of the 181 (77.7%) associations from our TIME germline discovery pipeline, the majority of variants were detected as *cis* associations (96.1%), aside from 7 (3.9%) *trans* associations (**Figure S4C).** Ten cancer relevant TIME-SNPs (4.6%) affected protein-coding regions **(Figure S4D).** In the case of *CTSS, HLA-B, TAP2, HLA-A, CTLA-4* and *APOE*, missense variants in coding regions were associated with expression differences. In addition, missense variants in *AP5B1* and *ERAP1* were associated with expression differences in *CTSW* and *ERAP2,* respectively **(Figure S4E)** and missense variants in *EGFL8* and *CCHCR1* were associated with differences in the MHC II signature described by Sayaman et al.

**Figure 3:**
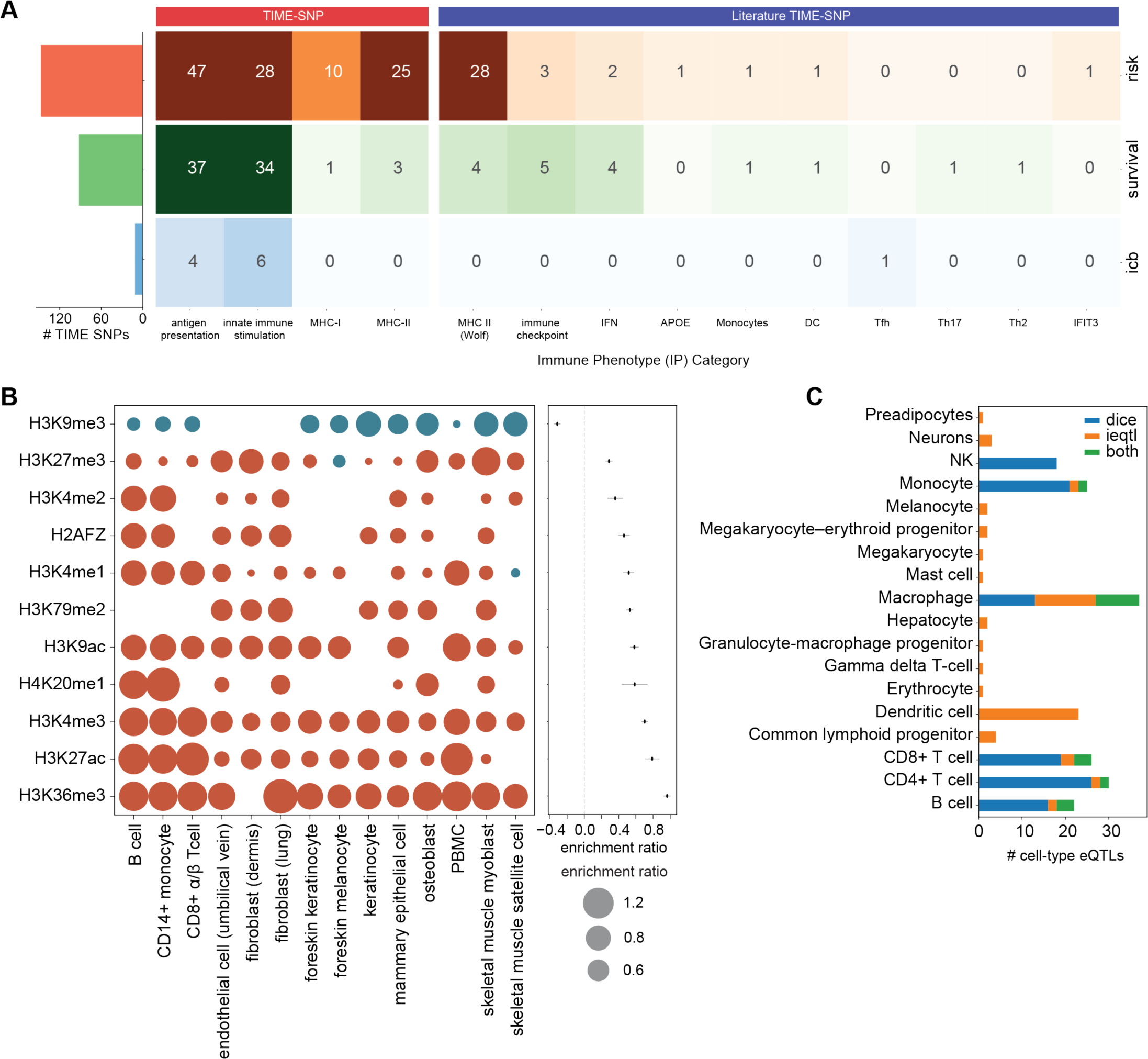
Identification and characterization of TIME-SNPs related to cancer outcomes. **(A)** Clustermap of cancer relevant associations with barplot of TIME-SNPs implicated in cancer risk, survival and immunotherapy response. **(B)** Mean enrichment ratio of immune microenvironment variants in histone marks with corresponding enrichment ratios in specific cell types. **(C)** Barplot of cell-type specific TIME-SNPs implicated by DICE and ieQTL analysis.

As the majority of TIME-SNPs fell within non-coding genomic regions, we evaluated their potential to affect regulatory sites. Histone marks provide important information about regulation of chromatin architecture and accessibility of DNA sequence for transcription^59^. Regions harboring TIME-SNPs were strongly enriched in H3K27ac, H3K36me3 and H3K4me3 histone marks and depleted in H3K9me3 histone marks **(Figure 3B).**^60^ H3K27ac is a known marker of active enhancers and H3K4me3 is usually enriched at promoters near transcription start sites^61, 62^ suggesting some TIME-SNPs are broadly associated with transcription while others may be gene specific. Coincidentally, TIME-SNPs were depleted in repressive H3K9me3 marks^63^. In addition to overall enrichment in markers of active transcription, enrichment in particular histone marks was more pronounced in certain immune cell types. The strongest enrichment in *cis*-regulatory elements was seen in PBMCs, CD14+ monocytes, B cells and CD8+ T cells **(Supplementary Table 9)**.

To obtain more information about TIME-SNP cell-type specific effects, we evaluated whether TIME-SNP association with gene expression in TCGA was dependent on immune cell infiltration level or corresponded to known immune eQTLs in DICE **(Methods)**. Of the 219 variants, 37 were macrophage cell-type eQTLs, 32 were CD4+ T cell-type eQTLs, 26 were CD8+ T cell-type eQTLs and 22 were B cell-type eQTLs **(Figure 3C**, **Supplementary Table 10)**. Comparing myeloid-specific eQTLs to lymphoid-specific eQTLs, variants associated with *TAP2, CTLA-4, FCGR3B, ERAP2* and *DBNDD1* were myeloid-specific. Looking specifically at cell types implicated by risk, survival and immunotherapy response, macrophages, CD4+ and CD8+ T cells were the cell types with the most associations **(Figures S4F)**. These findings reveal a subset of TIME-SNPs that specifically modify the activity of genes in immune rather than tumor cells and indicate cell types that may contribute to cancer risk, progression and immunotherapy response.

### TIME-SNPs underlying antigen presentation stratify melanoma and prostate cancer risk

We further analyzed genes with TIME-SNP cancer risk associations in the UK Biobank. At a strict FDR threshold (FDR < 0.05), 57 TIME-SNPs were associated with cancer risk, 28 of which were associated with genes critical for antigen presentation, such as *CTSS, ERAP1, CTSW, ERAP2* and MHC Class I and II genes. Using a less stringent FDR threshold (FDR < 0.20), we identified 119 TIME-SNPs associated with cancer risk, including 65 variants associated with genes critical for antigen presentation.

We noted an EP300 variant weakly associated with risk of neoplasms of digestive organs (FDR < 17%). *EP300* can potentiate MHC Class I antigen presentation^64^ and *EP300* somatic mutations have been reported in multiple types of cancer. Downregulation of *EP300* has been associated with higher anti-tumor immunity and suggesting that *EP300* inhibition could potentiate immune checkpoint blockade^65, 66^. However, in work by Kruper et al., higher EP300 expression was associated with more rapid tumor cell proliferation and sensitivity to ICB, while lower activity conferred resistance, suggesting instead that EP300 loss may potentiate immune evasion^64^. In our case, the variant allele was associated both with increased expression in the TCGA and with higher cancer risk in the UK Biobank, suggesting that more rapid growth could outweigh the benefit of immune evasion for early digestive tumors^65^. Cell-type interaction analysis did not reveal *EP300* variant interaction with any specific immune cell types.

To further validate antigen presentation TIME-SNPs as *bona fide* cancer risk SNPs, we asked whether a polygenic risk score (PRS) using antigen presentation TIME-SNPs would generalize to independent cancer cohorts. A TIME-SNP melanoma PRS restricted to associations with antigen presentation genes and the MHC Class II signature described in Sayaman et al. stratified patients with high vs low risk in the UK Biobank **(Figure 4A)** and validated in an independent cohort of 3029 melanoma cases and controls for UT MD Anderson^67^. Although the difference in PRS score distributions for cases and controls was small **(Figure 4B),** the odds of melanoma were significantly different in the top and bottom 10th quantile in the validation cohort **(Figure 4C).** The MHC II Signature described in Sayaman et al. was strongly correlated with MHC class II genes, such as *HLA-DQB1,* but also weakly correlated with MHC Class I gene expression **(Figure S5).**

**Figure 4:**
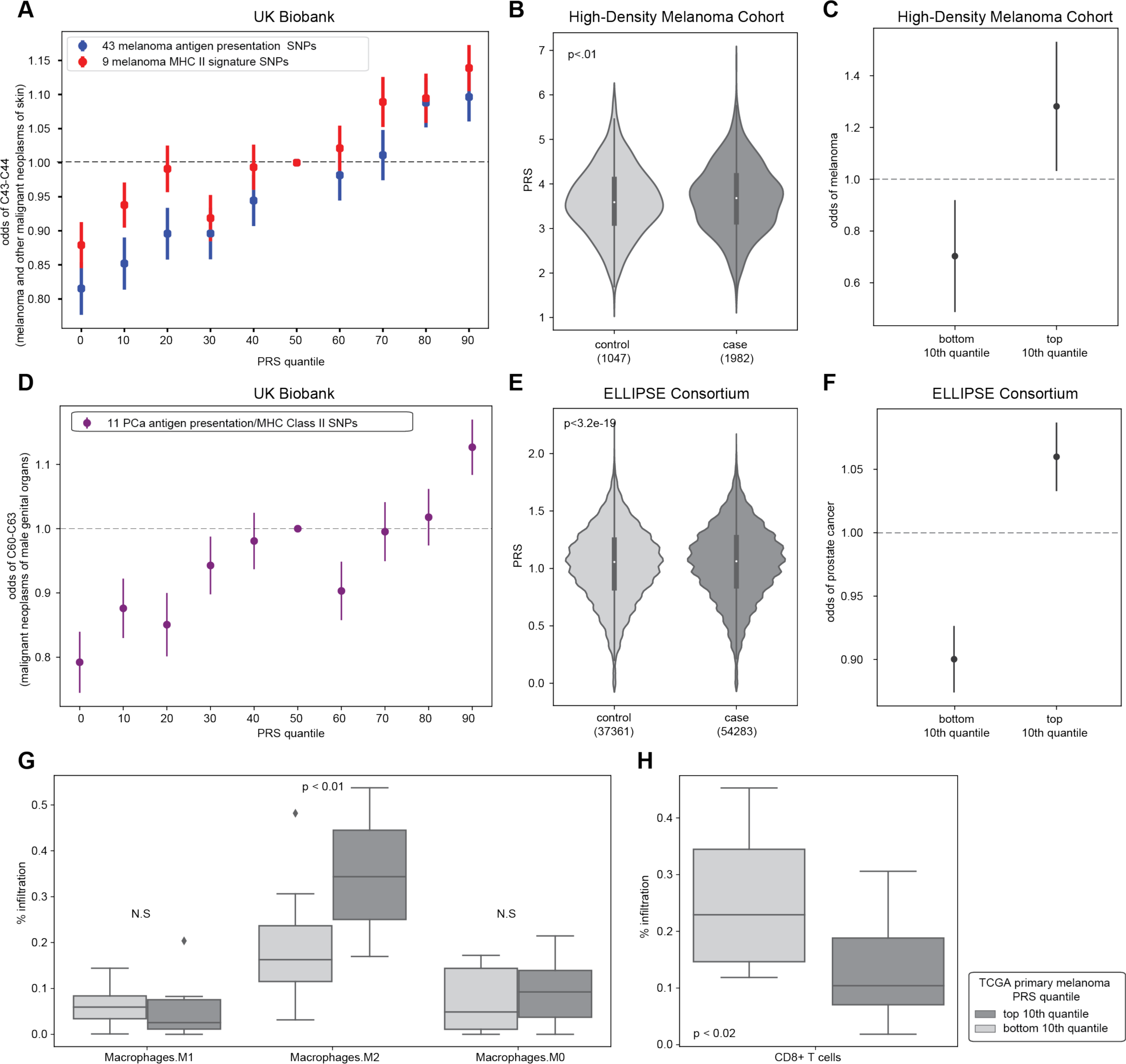
TIME-SNPs underlying antigen presentation stratify melanoma and prostate cancer risk. **(A)** UK Biobank quantile plot of polygenic risk score (PRS) constructed from antigen presentation gene TIME-SNPs and Sayaman et al. MHC II signature TIME-SNPs associated with melanoma risk. **(B)** Violinplot of PRS constructed from antigen presentation genes and MHC II signature TIME-SNPs in melanoma cases and controls in High Density melanoma cohort. **(C)** Odds of melanoma risk among individuals in the top and bottom 10th quantile of PRS in ELLIPSE consortium. **(D)** UK Biobank quantile plot of PRS constructed from antigen presentation gene TIME-SNPs and Sayaman et al. MHC II signature TIME-SNPs associated with prostate cancer risk. **(E)** Violinplot of PRS constructed from antigen presentation genes and MHC II signature TIME-SNPs in prostate cancer cases and controls in ELLIPSE Consortium. **(F)** Odds of prostate cancer risk among individuals in the top and bottom 10th quantile of PRS in ELLIPSE consortium **(G)** Boxplot of macrophage infiltration in primary TCGA SKCM (melanoma) in top and bottom 10th quantile of melanoma antigen presentation TIME- SNP PRS. **(H)** Boxplot of CD8+ T cell infiltration in TCGA SKCM (melanoma) in top and bottom 10th quantile of melanoma antigen presentation TIME-SNP PRS.

Eleven variants were associated both with prostate cancer risk in the UK Biobank (FDR < 0.2) and with the MHC II signature or antigen presentation genes, with 1 SNP in common with the melanoma PRS. A prostate cancer TIME-SNP PRS constructed from these in the UKBB stratified patients with high vs low risk **(Figure 4D)** and validated in an independent cohort of 91,644 prostate cancer cases and controls for ELLIPSE Consortium^68^. Although the difference in PRS score distributions for cases and controls was small in the validation cohort **(Figure 4E),** the odds of prostate cancer were significantly different in the top and bottom 10th quantile **(Figure 4F).**

Notably, 14 of the melanoma PRS TIME-SNPs and 4 of the prostate cancer PRS TIME-SNPs were DICE eQTLs. Of the melanoma PRS TIME-SNPs, 10 were macrophage cell-type eQTLs and 10 were CD8+ T cell-type eQTLs. This suggested that the PRS might be related to cancer risk through modulation of the inflammatory landscape. Indeed, in the TCGA, tumors in the upper 10th quantile of the melanoma TIME-SNP PRS had higher levels of infiltration by pro-tumor inflammatory M2-like **(Figure 4G),** but not M0 or M1-like macrophages. Promotion of an inflammatory pro-tumor environment was also correlated with decreased CD8+ T cell infiltration **(Figure 4H).**

### Variants underlying immune evasion are associated with cancer survival

We revisited the 92 significant associations and 87 unique variants implicated via Kaplan-Meier Analysis (FDR < 0.05) using the burden of SNPs in each of the categories shown in Figure 3A by performing a within tumor type Cox Proportional-Hazards analysis with covariates including age of diagnosis, gender, and stage of cancer, requiring associations to pass an FDR < 0.05. We found a significant association of immune checkpoint variant burden with overall survival in lung adenocarcinoma (FDR < 0.05) (**Figure 5A, S6A).** In the case of Th17 signature, Th2 signature, and dendritic cell signatures, we only had 1 variant per category. Nonetheless, these variants were significantly associated with progression-free survival (FDR < 0.05) (**Figure 5B, S6B)**. The variant associated with Th17 was an *IL17RA* intronic variant. We observed worse survival (**Figure 5C)** in liver and hepatocellular carcinoma patients with this variant that increased *IL17RA* expression in TCGA (**Figure 5D).** Indeed, this confirms reports that IL-17 is associated with worse survival in hepatocellular carcinoma through the potential pro-inflammatory role of IL-17^69, 70^.

**Figure 5:**
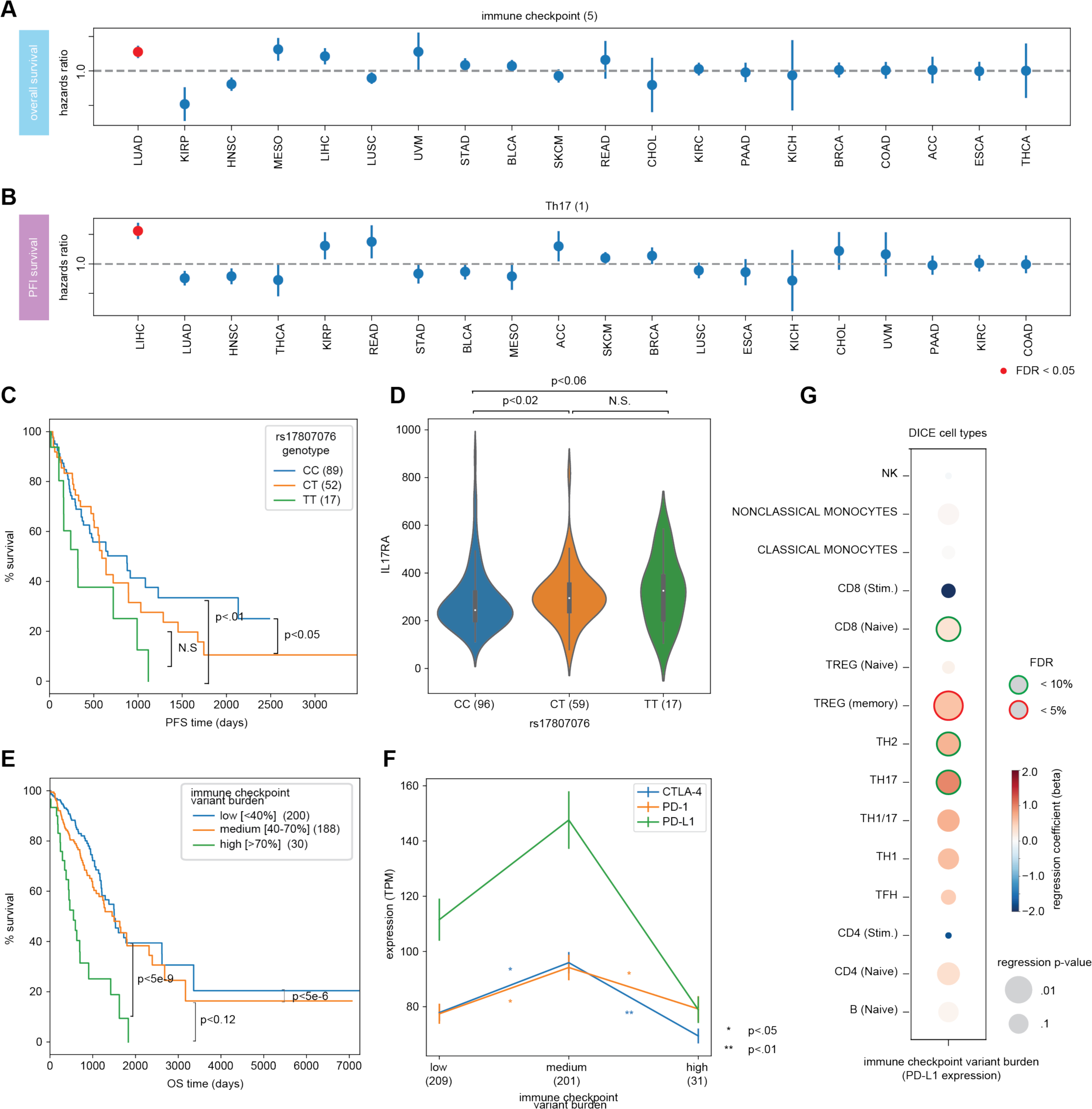
Variants underlying immune evasion are associated with cancer survival. **(A)** Cox Proportional- Hazards odds ratios for immune checkpoint burden score (constructed from 5 immune checkpoint variants) with overall survival separated by TCGA cancer type. **(B)** Cox Proportional-Hazards odds ratios for rs17807076 and progression-free survival separated by TCGA cancer type. **(C)** Progression-free survival Kaplan-Meier curve based on rs17807076 genotype in TCGA LIHC. **(D)** Violin plot of rs17807076 genotype and bulk RNA-seq expression of *IL17RA* in TCGA LIHC. **(E)** Overall survival Kaplan-Meier curve based on burden of 5 immune checkpoint variants (immune checkpoint variant burden) in TCGA LUAD. **(F)** Average expression of immune checkpoint molecules, *PD-L1, PD-1* and *CTLA-4*, stratified by immune checkpoint variant burden **(G)** Association of *PD-L1* expression with immune checkpoint variant burden in 15 immune cell types from DICE.

We also found the burden of immune checkpoint variants to be significantly associated with overall survival in lung adenocarcinoma (**Figure 5E).** Three variants in the burden score were associated with *PD-L1* and 2 with *CTLA-4*. Two *PD-L1* variants were in regions associated with the gene promoter while 1 was in a region associated with H3K27ac marks, suggesting variants may modify transcription of the genes **(Figure S6C).** Interestingly, while we did not see correlation of the burden score with *PD-L1* expression in bulk tumor RNA, we observed weak correlation with expression of *PD-1* and *CTLA-4* **(Figure 5F)**. Individuals with mid-level burden had increased expression of immune checkpoints and worse survival compared to individuals with a lower burden of variants, consistent with the immuno-inhibitory roles of these molecules. However, individuals with high immune checkpoint burden had significantly decreased immune checkpoint expression. This observation could point to an adaptive response of the tumor immune microenvironment or even potential non-genetic compensatory mechanisms that have developed to avoid high immune checkpoint expression throughout life. We investigated correlation of the burden score with tumor mutation burden (TMB) in lung cancers but found no significant correlation **(Figure S6D)**. Since immune checkpoint molecules have very cell-type specific expression, we investigated correlation of the burden score with expression of *PD-L1* specifically in immune cell types (**Figure 5G)** and indeed found positive correlation in immune cell subsets, except for stimulated CD8+ and CD4+ T cells. These results confirm that TIME-SNPs have cell-type effects, as we observed stronger effects of TIME-SNPs on immune cell specific expression compared to bulk RNA expression in the TCGA.

### TIME-SNPs implicate targets for modulating immune responses

To evaluate whether cancer relevant TIME-SNPs implicate aspects of tumor immunity that could serve as an entry point for immunotherapy, we further analyzed the 11 variants associated with ICB response, and their associated IP components **(Figure 6A)**. We confirmed that TIME-SNP genotypes were similarly distributed and free of artifacts across all cohorts by PCA analysis **(Figure S7A).** The direction of effect of variants associated with responder status was mostly consistent across cohorts, though there were some differences observed with the *PSMD11* variant, rs28459155 (**Figure 6B)**. This variant was associated with lower odds of being a responder in Miao et al. and Hugo et al. but higher odds of being a responder in Van Allen et al., Snyder et al. and Riaz et al. As a comparison to current ICB biomarkers, we also evaluated association of tumor mutation burden (TMB) and expression levels of *PD-L1, PD-1,* and *CTLA-4* with responder status and found no significant associations **(Figure 6B)**. We ran associations with the 11 variants and *TMB, PD-L1, PD-1,* and *CTLA-4* to determine if any variants were associated with these previously researched biomarkers. We observed an association between TMB and *ERAP1* variant rs27765 in Hugo et al. (**Figure S7B, Figure S7C).**

**Figure 6:**
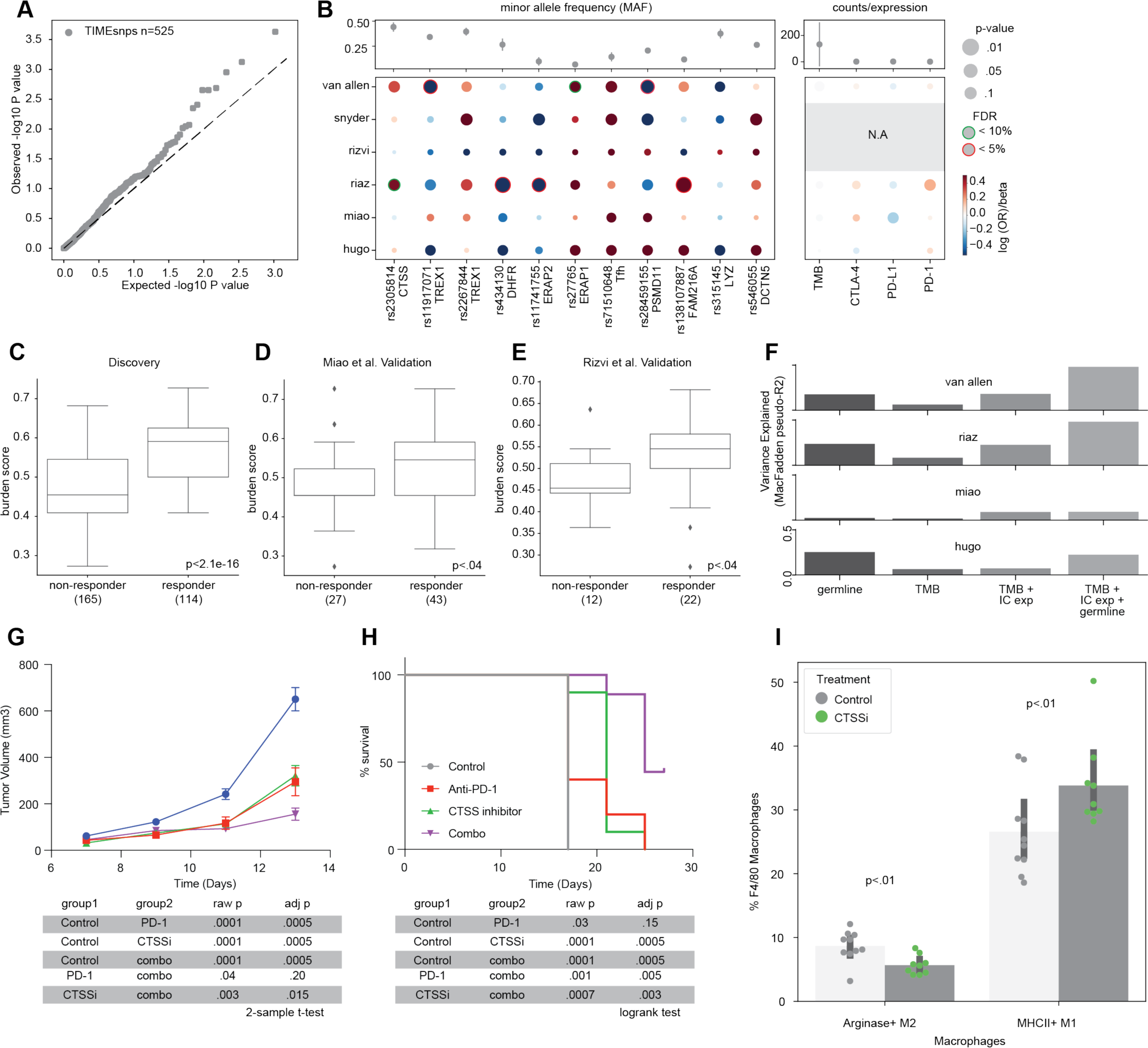
TIME-SNPs implicate targets for modulating immune responses. **(A)** Q-Q plot of METAL meta- analysis sample-size weighted association with immunotherapy response in Riaz et al., Snyder et al., Hugo et al. and Van Allen et al. melanoma discovery cohort. **(B)** Grid plot of log odds ratio of variants with responder status in 6 ICB cohorts with beta coefficients of classic ICB biomarkers (TMB, *PD-L1, PD-1, CTLA-4*) association with responder status. **(C)** Boxplot of burden score constructed from 11 significant ICB variants (FDR < 0.5) in discovery melanoma cohort. **(D)** Boxplot of burden score constructed from 11 significant ICB variants (FDR < 0.5) in Miao et al validation cohort. **(E).** Boxplot of burden score constructed from 11 significant ICB variants (FDR < 0.5) in Rizvi et al validation cohort. **(F)** Pseudo-R^2^ (variance explained) with germline burden score only, TMB only, expression of *PD-L1, PD-1* and *CTLA-4*, and germline burden score with TMB and expression of *PD-L1, PD-1* and *CTLA-4*. **(G)** Tumor growth curve for C57BL/6 mice implanted with MC38 treated with anti- PD-1, anti-CTSS, and combination of anti-PD-1 and anti-CTSS. (**H)** Survival curve for C57BL/6 mice implanted with MC38 treated with anti-PD-1, anti-CTSS, and combination of anti-PD-1 and anti-CTSS. **(I)** Barplot of the proportion of F4/80 Macrophages that are Arginase^+^ M2 macrophages and MHCII^+^ M1 macrophages respectively for MC38 tumors treated with anti-CTSS compared to control.

We constructed a burden score with the response-associated allele for all 11 variants across the four melanoma ICB cohorts and confirmed that ICB responders had a significantly higher score compared to non- responders (**Figure 6C**). The association between higher burden score and ICB response remained significant in two additional independent validation cohorts from Miao et al. (renal cell carcinoma) and Rizvi et al. (non-small cell lung cancer) (**Figure 6D, 6E**). We repeated this analysis selecting 11 TIME-SNPs at random, matched for minor allele frequency, and found that the observed difference in burden score between responders and nonresponders was significantly larger than random in both discovery and validation sets (Figure S7D,E).

Although we had a small sample size, we built a polygenic risk score trained on the discovery melanoma cohort using PRSice and were able to achieve an AUC of at least 0.6 in Rizvi et al. and Miao et al. validation sets (Figure S7F). We then evaluated how much of the variation in responder status was explained by germline burden score, TMB, TMB plus immune checkpoint markers, and all of these markers together. Germline burden score explained some variance in responder status in all 6 cohorts (2.5-25.4%). In Van Allen et al., Hugo et al., and Riaz et al., germline burden score explained more variance than TMB and immune checkpoint markers. In Van Allen et al. and Riaz et al., the combination of all markers maximized the variance explained (**Figure 6F**).

Colocalization of gene expression and GWAS signals can point to putative causal disease-related genes. To determine if genes implicated by TIME-SNPs could reveal candidate immunotherapy targets, we analyzed the genes associated with the 11 variants. These genes once again emphasized a central role for antigen processing and presentation pathways (GO:0048002 antigen processing and presentation of peptide antigen [fold enrichment = 90.33, FDR < 0.0351], GO:0030333 antigen processing and presentation [fold enrichment = 82.46, FDR < 0.0011]). Examining gene expression available for 4 of the 6 cohorts, we noted that none of these genes were significantly differentially expressed between ICB responders and nonresponders (**Supplementary Table 11**). However, some TIME-SNPs were associated with high expression of an IP component gene and worse outcome, uncovering potential targets to validate. Two *TREX1* variants were associated with immunotherapy response; rs11917071 was associated with increased *TREX1* expression in Van Allen et al. and lower odds of being a responder (Figure S7G), and rs2267844 was associated with decreased *TREX1* expression in Hugo et al. and Miao et al. and higher odds of being a responder (Figure S7H). These results are consistent with findings that *TREX1* acts as an immunoinhibitor that prevents cGAS-STRING initiation, with inhibition of *TREX1* stimulating IFNγ signaling and autoimmunity, making it a potential immunomodulatory target^71, 72^. *PSMD11* variant rs28459155 was associated with lower odds of being a responder but increased expression of *PSMD11* (Figure S7I)*. PSMD11*, a proteosomal protein involved in ubiquitination, is associated with worse prognosis in pancreatic cancer^73^. Similarly, *CTSS* variant rs23058814 was associated with higher odds of being a responder and decreased *CTSS* expression in Van Allen et al. (Figure S7J). Increased *CTSS* expression has been linked to tumor progression in follicular lymphoma due to decreased CD8+ T cell recruitment^64^. In two separate mouse models, significant differences in *Ctss* expression was observed between immunotherapy responders and non-responders (Figure S7K,L). Furthermore, we observed increased M1 macrophage infiltration in individuals with *CTSS* variant in Hugo et al. (Figure S7M).

Based on these association data, we hypothesized the inhibition of *CTSS* would improve ICB response. To test this hypothesis, we treated mice implanted with MC38 tumors with a CTSS small molecule inhibitor. Mice treated with CTSS inhibitor had slowed tumor growth and better survival compared to control mice (**Figure 6G, 6H**). We also evaluated the interaction of CTSS inhibitor treatment with anti-PD-1. Mice treated with CTSS inhibitor or anti-PD-1 monotherapy had significantly decreased tumor growth and better survival compared to control mice. Additionally, tumor growth was further decreased in mice treated with the combination of anti-PD- 1 and CTSS inhibitor as compared to mice treated with anti-PD-1 or CTSS inhibitor alone. In the MC38 model, we observed an increase in infiltrating M1 macrophages and a decrease in M2 macrophages similar to findings from Hugo et al. (**Figure 6I**). These findings demonstrate that a focused screen for cancer relevant TIME- associated variants provides a fruitful strategy to reveal novel immunotherapy targets. Furthermore, the influence of *CTSS* inhibition on the myeloid landscape, identifies macrophages as potential cell type that may modulate immunotherapy response.

## Discussion

The success of immunotherapies has generated enthusiasm for using the human immune system as a weapon to eliminate cancers^74–77^. However the very existence of cancer indicates the failure of the immune system to control malignant cell populations throughout multiple stages of tumor development^4^. Here we studied genetic variants associated with interindividual differences in immune traits and the tumor immune microenvironment, reasoning that these variants could reveal the aspects of immunity most critical for the successful immune control of tumors. We focused on a subset of immune characteristics that showed evidence of SNP heritability in The Cancer Genome Atlas or were implicated in the literature and that were also associated with cancer outcomes including risk, survival and response to immune checkpoint inhibitors. Out of the 1084 SNP associations with 129 immune phenotype (IP) components, ultimately 219 TIME-SNPs associating with 65 IP components met these criteria. 104 of these SNPs were not in LD (R^2^ < 0.5) with genome-wide significant SNPs reported by cancer GWAS studies or Vanderbilt PheWAS catalog despite interacting with important clinical outcomes. In addition, we demonstrated that our approach could implicate putative targets to modify anti-tumor immunity; *TREX1* has previously been highlighted as a promising target^71^, and small molecule inhibition of *CTSS* resulted in slower tumor growth and longer survival of mice, with effects comparable to anti-PD-1. While several studies suggest that CTSS may have immune suppressive roles in follicular lymphoma^78–80^, this is the first time to the best of our knowledge that inhibition of this gene was shown to relieve immune suppression in solid tumors.

The immune system interacts with tumors throughout their development and treatment, both through tumor-promoting inflammation and immune-mediated elimination of cancerous cells^81, 82^. Cancer relevant SNP associations implicated IP components related to antigen presentation (MHC Class I and II, *CTSS*, *ERAP1*) and inflammation (Th17, Th2). Limited sample sizes and the inability to impute HLA region SNPs with sufficient quality in exome-only cohorts impeded the complete assessment of pleiotropy in the context of cancer relevance. Nonetheless, we found several SNPs linked to genes involved in antigen presentation and evasion of the adaptive immune response associated with multiple aspects of the tumor-immune interplay. More generally, we saw that IP components implicated in cancer risk were mainly those involved in both MHC Class I and Class II antigen presentation, while SNPs associated with prognosis pointed to genes that would support evasion of the MHC I CD8+ T cell axis including *PD-L1, CTLA-4* and Th17.

Adaptive immunity is crucial for the host anti-tumor immune responses. In cancer, downregulation of antigen presentation machinery is a frequent mechanism of immune evasion. MHC Class I and II genotypes shape mutational landscapes in cancer and inform immunotherapy response^16, 17, 20^. Interestingly, we found multiple germline variants associated with both MHC Class I and Class II genes were also associated with cancer risk. This included several antigen presentation pathway genes not directly encoding the MHC and located outside of the HLA region: *CTSS, CTSW, ERAP1, ERAP2,* and *TAP2. ERAP1* and *ERAP2* are endoplasmic reticulum peptidases that trim peptides before loading them onto MHC proteins^83, 84^. *ERAP1/ERAP2* polymorphisms have been associated with cervical cancer and autoimmunity^85–91^. *CTSS* is a cysteine protease critical for MHC Class II loading and is frequently mutated in follicular lymphoma. Its loss limits communication with CD4+ T follicular helper cells while inducing antigen diversification and activation of CD8+ T cells^78, 79^. *CTSW* is crucial for cytotoxicity and is expressed in specific immune cell types^79^. Interestingly, the involvement of MHC II and immune cell specific genes suggest that inter-individual variation in immune surveillance contributes to cancer risk. In further support of this, polygenic risk scores constructed from MHC II pathway associated germline variants were able to stratify melanoma and prostate cancer risk in multiple cohorts. These two tumor types fall at opposite ends of the spectrum of immune activity, with melanoma being one of the most immunotherapy responsive tumors, and prostate cancer being one of the least^92, 93^. Furthermore, MHC Class II expression is linked to ICB response in melanoma^94^. Although prostate cancer is considered immunologically “cold”, rare dramatic responses to immunotherapy have been documented^95^. MHC Class II is usually restricted to professional antigen presenting cells although prostate cancer cells have been shown to express MHC Class II. The Class II pathway is crucial for a prolonged anti-tumor response as it leads to sustained CD8+ T cell activation and leads to more complete tumor clearance. Due to low imputation quality, we were not able to assess the dependence of ICB responses on HLA region SNPs, however *CTSS* was both detected and validated as a determinant of response, suggesting MHC Class II could also underlie response to immunotherapy. Together with reports from multiple immune vaccines studies that responses were primarily driven by CD4+ T cells^96–99^, these findings place further emphasis on the central importance of MHC II for effective anti-tumor immune responses.

Innate immunity is the branch of the immune system that acts as the body’s first line of defense against microbial pathogens and cancer cells, and involves cells originating in the bone marrow that carry non- polymorphic receptors. Cells of the innate branch of the immune system such as macrophages and dendritic cells play a pivotal role in the tumor microenvironment creating a hostile pro-inflammatory environment, suppressing T cells, promoting angiogenesis, and initiating lymphangiogenesis. *TREX1*, *TLR2*, *VAMP3* and *APOE* were among innate immune genes implicated by our screen. In a recent study it was shown that mice with one missense mutation (D18N) in the TREX1 gene were protected from tumor growth, and that protection was associated with a reduced expression of *PD-1* in T lymphocytes (aka less exhaustion)^100^. This study supports the view that a missense mutation of TREX1 leads to enhanced tumor T cell immunity. Using RNA-seq analysis, we found individuals with *TREX1-*reducing variants had higher odds of being ICB responders. Arguably, TREX1 inhibition could be regarded as a viable anticancer immunotherapeutic strategy.

In general, the innate compartment was implicated mostly through cell type eQTL analysis. Variants associated with *ERAP1, CTSW, DCTN5, MICA, BTN3A2* and MHC Class I and II were cell for cells in both innate and adaptive subsets. *CTSW* was a cell-type eQTL for macrophages and dendritic cells, as well as CD8+ T cells and B cells. Although *CTSW* has strong immune expression in cytotoxic cells, such as NK cells and CD8+ T cells, previous literature has found *CTSW* to be prognostic and correlated with infiltration of both innate and adaptive cell types^101^. rs61802301, associated with *FCGR2B,* was specifically an cell-type eQTL for B cells; *FCGR2B* is crucial for B cell regulation^102^. However, variants associated with *TAP2, CTLA-4, FCGR3B, ERAP2,* and DBNDD1 were cell-type eQTLs primarily for myeloid cells, specifically antigen-presenting cells such as macrophages and dendritic cells. *CTLA-4* is expressed on dendritic cells and induces IFNγ production by dendritic cells^103^. We also observed that targeting *CTSS* resulted in a higher abundance of inflammatory M1 macrophages versus suppressive M2 macrophages, consistent with suppressor myeloid cells impairing effective anti-tumor immunity and ICB response^104, 105^. The abundance of innate cell cell-type eQTLs affecting adaptive genes may suggest that a disconnect between innate and adaptive function forms the basis of dysregulated adaptive immunity in the tumor immune microenvironment.

Notably a SNP-based burden score reproducibly correlated to ICB response across multiple cohorts with melanoma, non-small cell lung cancer (NSCLC) and kidney cancer (RCC). Furthermore, this burden score compared favorably with other popular measures such as tumor mutation burden (TMB) and checkpoint gene expression for predicting binary response category. In renal cell carcinoma the link between tumor mutation burden (TMB) and ICB response is not clear, in contrast to high TMB diseases like melanoma and NSCLC where higher TMB is associated with better responses^55, 106–108^. Possibly, in a setting with low TMB such as in RCC, host genetics have more value as prognostic biomarkers. In the future, germline determinants of the TIME could be integrated into predictors alongside other characteristics of the tumor immune microenvironment that have been found to inform immune response such as TMB, PD-L1 positivity, the number and quality of T cells^109^*, IFN-ɣ* response, cytotoxicity scores, T cell activation and T cell exhaustion signatures^53–55, 110–117^. Some of these factors require profiling of tumor RNA which is less commonly performed in clinical settings. If germline variants could serve as a proxy for characteristics of the TIME that otherwise require more complex molecular profiling, they could provide an avenue for more cost effective tools for the clinic.

A potential limitation of the applicability of our approach is that discovery of TIME-SNPs is dependent on the availability of paired genomic and transcriptomic data from tumors, which is currently available only for a few cohorts. Effect sizes associating genetic variants with cellular phenotypes are likely to be larger than those linking genetic variants to diseases^118–120^, however the number of associations detected may still be limited by available sample sizes and the limited population diversity thereof. We were able to impute a subset of our SNPs into existing immune checkpoint blockade study cohorts that had only exome sequencing, but others falling outside of exonic regions could not be analyzed in this context. Studies focused on tumor exomes and transcriptomes could include genome-wide SNP profiling via arrays or low pass whole genome sequencing to allow more effective integration into future studies of germline genomic variation.

## Conclusion

Interindividual differences encoded by germline variation contribute to considerable population variation in heritable traits, including in the immune system. Our analysis highlights the utility of studying this variation in the context of the tumor immune microenvironment to understand the aspects of immunity most germane to tumor development and uncovering novel immunotherapy targets. SNPs associated with variable characteristics of the TIME highlighted the major factors underlying anti-tumor immune responses. TIME-SNPs associated with cancer risk or response to immunotherapy merit further investigation as possible entry points for future efforts to engineer the tumor immune microenvironment.

## Acknowledgements

This work was supported by Emerging Leader Award from The Mark Foundation for Cancer Research, grant #18-022-ELA to HC, NIH grant R01CA220009 to HC and MZ, U01CA196406 to OH, and 1F30CA247168-01 and T32CA067754 to MP. UK Biobank data was retrieved under project ID 37671.

The results shown here are in large part based upon data generated by the TCGA Research Network: https://www.cancer.gov/tcga and Genotype-Tissue Expression (GTEx) Project: https://gtexportal.org/home/. GTEx was supported by the Common Fund of the Office of the Director of the National Institutes of Health, and by NCI, NHGRI, NHLBI, NIDA, NIMH, and NINDS. The data used for the analyses described in this manuscript were obtained from the GTEx Portal on 10/10/20. This research has been conducted using the UK Biobank Resource under project ID 37671. Cancer risk validation cohorts were obtained from dbgap accessions: phs000187.v1.p1 and phs001125.v1.p1. Research support to collect data for phs000187.v1.p1 and develop an application to support phs000187.v1.p1 was provided by 3P50CA093459, 5P50CA097007, 5R01ES011740, and 5R01CA133996. Funding for the meta-analysis in phs001125.v1.p1 is provided by NIH grant U19CA148537. We would like to acknowledge the NCRN nurses and Consultants for their work in the UKGPCS study. We thank all the patients who took part in this study. This work was supported by Cancer Research UK (grant numbers C5047/A7357, C1287/A10118, C1287/A5260, C5047/A3354, C5047/A10692, C16913/A6135 and C16913/A6835). We would also like to thank the following for funding support: Prostate Research Campaign UK (now Prostate Cancer UK), The Institute of Cancer Research and The Everyman Campaign, The National Cancer Research Network UK, The National Cancer Research Institute (NCRI) UK. We are grateful for support of NIHR funding to the NIHR Biomedical Research Centre at The Institute of Cancer Research and The Royal Marsden NHS Foundation Trust. The MEC was supported by NIH grants CA63464, CA54281 and CA098758. For Rizvi *et al.* non-small cell lung cancer immunotherapy analysis, we used dbGaP data from accession phs000980.v1.p1. We thank the members of the Thoracic Oncology Service and the Chan and Wolchok labs at MSKCC for helpful discussions. We thank the Immune Monitoring Core at MSKCC, including L. Caro, R. Ramsawak, and Z. Mu, for exceptional support with processing and banking peripheral blood lymphocytes. We thank P. Worrell and E. Brzostowski for help in identifying tumor specimens for analysis. We thank A. Viale for superb technical assistance. We thank D. Philips, M. van Buuren, and M. Toebes for help performing the combinatorial coding screens. The data presented in this paper are tabulated in the main paper and in the supplementary materials. This work was supported by the Geoffrey Beene Cancer Research Center (MDH, NAR, TAC, JDW, AS), the Society for Memorial Sloan Kettering Cancer Center (MDH), Lung Cancer Research Foundation (WL), Frederick Adler Chair Fund (TAC), The One Ball Matt Memorial Golf Tournament (EBG), Queen Wilhelmina Cancer Research Award (TNS), The STARR Foundation (TAC, JDW), the Ludwig Trust (JDW), and a Stand Up To Cancer-Cancer Research Institute Cancer Immunology Translational Cancer Research Grant (JDW, TNS, TAC). Stand Up To Cancer is a program of the Entertainment Industry Foundation administered by the American Association for Cancer Research. For Snyder et al. melanoma immunotherapy analysis, we used dbGaP data from accession phs001041.v1.p1. We thank Martin Miller at Memorial Sloan Kettering Cancer Center (MSKCC) for his assistance with the NetMHC server, Agnes Viale and Kety Huberman at the MSKCC Genomics Core, Annamalai Selvakumar and Alice Yeh at the MSKCC HLA typing laboratory for their technical assistance, and John Khoury for assistance in chart review. For Miao et al renal cell carcinoma immunotherapy analysis, we used dbGap data from accession phs001493.v2.p1. This study was supported by an AACR KureIt grant. Hugo *et al.* melanoma samples were acquired from SRA using accession numbers SRP067938 and SRP090294. Riaz *et al.* melanoma samples were acquired from SRA using accession number SRP095809. For Van Allen *et al.* melanoma sample, data was acquired from dbgap accession phs000452.v2.p1.

## Author Contributions

M.P. and H.C. conceived the work and designed and analyzed the experiments; W.T. and C.F. assisted in statistical analyses; M.P. and H.C. wrote the paper with assistance from P.V., W.T., M.Z., J.M., S.P., O.H., C.F., G.M., C.P.D and A.C.; A.C. and G.M. advised on HLA region analysis; J.T. performed optimization of risk score analysis; B.J.S. and C.G.C. performed DICE annotation under supervision of P.V.; T.S. conducted DESeq2 analysis; H.K. conducted UK Biobank PheWAS analysis; S.C. and R.M.S. performed UK Biobank preprocessing used in PheWAS analysis; V.W. and S.G., performed MC38 mouse experimental validation and analyzed results; C.P.D, E.G. and G.M. provided M4 mouse validation results; S.G. assisted in epigenetic analysis; D.K. assisted in ieQTL analysis

## Resource Availability

Further information and requests for resources should be directed to and will be fulfilled by the Lead Contact, Hannah Carter.

## Data Availability

The data that support the findings of this study are available on request from the corresponding authors H.C. and M.P. The data are not publicly available as they were accessed through dbGaP applications and are controlled data that could lead to reidentification of individuals. For Data Access to processed genotyping, transcriptomic and mutation data, contact corresponding authors with proof of access to dbGaP studies.

## Code Availability

All code used for analysis and figure generation are available at https://github.com/meghanasp21/TIMEgermline.

## Supplementary Tables

**Supplementary Table 1:** Description of IP components

**Supplementary Table 2:** Genome-wide Complex Trait Analysis (GCTA) results

**Supplementary Table 3:** Significant TIME Associations from GWAS Analysis of 157 IP Components

**Supplementary Table 4:** PCA loadings from analysis of 157 IP components

**Supplementary Table 5:** Significant TIME Associations from Literature

**Supplementary Table 6:** UK Biobank PheWAS Results of TIME-SNPs

**Supplementary Table 7:** Kaplan-Meier Survival Analysis of TIME-SNPs

**Supplementary Table 8:** METAL Immune-Checkpoint Blockade (ICB) association analysis results

**Supplementary Table 9:** GREGOR histone mark enrichment analysis of TIME-SNPs

**Supplementary Table 10:** Significant TIME-SNP cell-type eQTLs

**Supplementary Table 11:** DESeq2 analysis of ICB-variant TIME genes

## Method Details

### TCGA Subject Details

The Cancer Genome Atlas (TCGA) consists of tumor and matched normal samples for over 11,000 patients. The Genomic Data Commons (GDC) legacy archive contains germline data for 11,542 samples from 10,875 unique individuals. Samples with TCGA project IDs: DLBC, LAML, THYM were excluded as they represent cancers derived from immune cells. Pairs of individuals with estimated KING kinship coefficient > 0.177, which represents first-degree relatedness were excluded. TCGA individuals were consented for general research use and no attempts were made to reidentify or contact subjects. Both females and males were included, and sex and individual age were included as covariates. Experimenters were not blinded and randomization of subjects was not relevant to the study.

### TCGA Genotypes

Normal (non-tumor) level 2 genotype calls generated from Affymetrix SNP6.0 array intensities using BIRDSUITE (RRID: SCR_001794) software^121^ were retrieved from TCGA GDC Legacy Portal (accession date: 04/26/2019). In these files, each of 906600 SNPs was annotated with an allele count (0=AA, 1=AB, 2=BB, - 1=missing) and confidence score between 0 and 1. Genotypes with a score larger than 0.1 (error rate > 10%) were set to missing and data were reformatted for PLINK (RRID:SCR_001757)^30^. We discarded 322 SNPs with probe names that did not match the hg19 UCSC Genome Browser (RRID:SCR_005780) Affymetrix track (track: SNP/CNV Arrays, table:snpArrayAffy6). Allele counts were converted to alleles using the definitions in metadata distributed with Affymetrix SNP 6.0 Array Documentation and negative strand genotypes were flipped to the positive strand using PLINK (PLINK, RRID:SCR_001757).

Pre-imputation processing of autosomal and X chromosome genotypes consisted of the following steps:

1. SNPs with call rate <90% were removed
2. SNPs with minor allele frequency (MAF) <1% were removed
3. Individuals with genotype coverage <90% were removed
4. Individuals with conflicting gender assignments were flagged
5. Heterozygous haploid SNPs were set to missing.

After applying these filters, the remaining 800644 autosomal and 32809 X chromosome SNPs were input to the secure Michigan Imputation Server^122^. SNPs were imputed with Minimac3/Minimac4 and European HRC Version r1.1 2016 reference with Eaglev2.3 phasing.

Post-imputation processing of genotypes included:

1. SNPs with MAF <1% were removed
2. Autosomal SNPs with Hardy-Weinberg Equilibrium <1e-9 were removed
3. Individuals with high heterozygosity rates (>3 SDs of mean) were removed
4. Pairs of individuals with kinship coefficient > 0.177 (first-degree relatedness) were removed

Rsq values from INFO files were extracted to annotate genotyping quality. The final genotyping data included 8217 individuals and 7,884,718 variants.

### TCGA Population Stratification

Ancestry filtering was applied using two techniques: (1) k-means clustering and (2) outlier identification. For k-means clustering, TCGA and HapMap Phase III populations were combined. HapMap Phase III genotypes were obtained from the NCBI HapMap ftp site and lifted to hg19 using the liftOver utility^123^. Genotypes were merged and reduced to a set of 33,675 independent SNPs determined previously^120, 123^ through linkage-based filtering using PLINK (RRID:SCR_001757). Pairwise identity-by-state (IBS) between all individuals was calculated and the resulting IBS matrix was used for PCA analysis. K-means clustering trained on HAPMAP Phase III separated individuals into the following groups: (1) TSI, CEU, (2) JPT, CHD, CHB, (3) MEX, (4) GIH, MKK, (6) YRI, ASW, LWK. This trained model was used to predict groups in TCGA. Cluster (1) were identified as European individuals.

We ran the aberrant R package v1.0 with lambda 20 for outlier identification^124^. Intersection of k-means clustered individuals and non-outlier individuals from outlier identification analysis was used to include TCGA individuals in the European ancestry discovery cohort.

### TCGA Phenotype Data

PanCanAtlas RNA data from GDC PanCanAtlas Publications Supplemental Data (https://gdc.cancer.gov/about-data/publications/pancanatlas) was downloaded (access date: 10/14/19). Only primary tumors (barcode: 01A/01B/01C) were considered in our analysis. Corresponding clinical metadata were obtained from the GDC Portal (https://tcga-data.nci.nih.gov/docs/publications/tcga/).

The following phenotypes were extracted or generated from RNA-seq data:

1. **Immunomodulators:** 436 genes used to define immune states from Thorsson et al.
2. **Immune checkpoint molecules:** 78 immune checkpoint stimulatory and inhibitory molecules from Thorsson et al.
3. **Antigen Presentation:** 231 antigen presentation genes from Gene Ontology [GO_REF:0000022]
4. **Immune cell markers:** 60 immune cell type markers from Danaher et al.
5. ***IFN-ɣ*:** *IFN-ɣ* genes retrieved from Biocarta [Systematic Name: M18933]
6. ***TGF-β*:** *TGF-β* genes retrieved from Biocarta [Systematic Name: M22085)
7. **Immune states:** Individual level scores for 6 immune states [wound healing, *IFN-ɣ* dominant, inflammatory, lymphocyte depleted, immunologically quiet, and *TGF-β* dominant] from Thorsson et al.
8. **Immune infiltration levels:** 22 relative immune infiltration estimates from CIBERSORTx^125^ using LM22 signature matrix.

Phenotypes with greater than 10% zero values were excluded and rank-based inverse normal transformation (**Figure S1)** was applied to each tissue type using **Eqn 1**^126^.

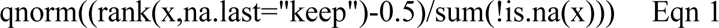

A total of 733 phenotypes remained for preliminary analyses.

### Heritability Estimates

Heritability estimates were calculated with the genomic-relatedness-based restricted maximum-likelihood (GREML) approach implemented in GCTA (Genome-wide Complex Trait Analysis)^127, 128^. Genetic relationship matrices (GRMs) which measure genetic similarity of unrelated individuals (GRM <0.05) were constructed for the autosomal and X chromosomes for the European cohort. Benjamini-Hochberg false discovery rates (FDR) were calculated using statsmodels^129^. Immune traits were considered sufficiently heritable if the V(g)/V(p) value was > 0.05 using the full GRM.

As highly polymorphic regions such as HLA and KIR gene regions can inflate heritability estimates, we conducted a 2-state GCTA analysis with separate GRMs for HLA/KIR regions (HLA chr6:28,477,797- 33,448,354, KIR chr19:55,228,188-55,383,188) and with the rest of the genome excluding HLA/KIR regions. Age and sex were included as covariates. An FDR < 0.05 was used to identify SNP-heritable IP components from 2-state analysis. If an IP component had high SNP heritability using the HLA/KIR GRM, a conditional GWAS analysis was conducted; otherwise, a standard GWAS analysis with Bonferroni-corrected suggestive p-value threshold was conducted. Ultimately, 140 IP components outside of the HLA/KIR regions and 17 IP components within the HLA/KIR regions were identified.

### Principal Component Analysis

157 SNP-heritable components were analyzed using sklearn. IP component values were scaled by Sklearn Standard Scaler and used for principal component analysis (PCA). Ordinary least squares (OLS) regression was performed with 157 IP components and principal components, wherein the beta coefficient represents the degree of change in principal component for every unit change in IP component. P-values indicate whether a coefficient was significantly different from 0.

### GWAS Analysis

The GLM method in PLINK (RRID:SCR_001757) was used to conduct association analyses with IP components. All associations were adjusted for covariates of age, sex and the first ten principal components. Gene expression values, CIBERSORTx relative infiltration estimates, and immune state scores were inverse-rank normalized by tissue type to control for tissue-type expression effects. Significant associations were identified with the PLINK (RRID:SCR_001757) clumping method using the primary suggestive threshold corrected for the number of phenotypes tested^130^ (1x10-5/140) using a kb threshold of 500, and an R^2^ threshold of 0.5.

To determine if variants had been implicated in previous cancer GWAS studies, variants were input into the LDlink server (https://ldlink.nci.nih.gov/?tab=ldtrait) using parameters “EUR” population, an R^2^ threshold of and base pair window of 500kb^46, 47^. We also retrieved the Vanderbilt PheWAS catalog^48^ and any TIME-SNPs in high linkage disequilibrium (R^2^ > 0.5) with Vanderbilt PheWAS catalog cancer risk SNPs were included as cancer risk variants. Lastly, we assessed SNPs by PheWAS analysis in the UK Biobank (detailed below).

### Conditional HLA analysis

The PLINK (RRID:SCR_001757) GLM method was used to run stepwise conditional analysis for identification of independent HLA associations^34^. The most significant initial associations detected with HLA region phenotypes by standard GWAS analysis were incorporated as covariates in the subsequent round. Specifically, we re-ran the analysis with chromosome 6 variants including the most significant SNP (lowest p-value in the previous round) as a covariate. Analysis was conducted until no SNPs with Bonferroni-corrected p-value < (1x10^-5^/17) remained.

### HLA Allele Specific Expression

TCGA tumor specific RNA BAM files were downloaded from the GDC on 07/16/2019. The HLApers^131^ kallisto- based pipeline was used with gencode.v30 annotations^132^. Default parameters were used and the two alleles with the highest calculated expression were retained for each HLA gene if there were more than 2 alleles reported. The top 2 highest expressed HLA alleles for each gene were averaged for input into SNP analyses. If expression for at least two alleles was not calculated, expression was set as missing for the sample. Only primary samples (01A/01B/01B) were considered for analysis. Summed HLA allele specific expression was inverse-rank normalized by cancer type and used for downstream analyses.

### Literature TIME-SNPs

We compiled existing germline variants associated with the tumor immune microenvironment (TIME) or ICB response from the literature. We collected 14 studies with their descriptions below:

1. Kogan et al. *JCI* 2018: Discovery of *FGFR4* germline variant which enhances *STAT3* activity impeding CD8 T cell infiltration
2. Queirolo et al. *Front Immunol* 2017: Investigation of 6 *CTLA-4* SNVs in 173 metastatic melanoma patients with overall response and survival information
3. Uccellini et al. *J Transl Med* 2012: *IRF5* polymorphism was associated with non-response to adoptive therapy with TILs
4. Bedognetti et al. *Br J Cancer* 2013: *CXCR3* and *CCR5* genetic polymorphisms were evaluated for expression of respective ligands and TIL migration
5. Lim et al. *PNAS* 2018. Systematic identification of germline genetic polymorphisms associated xCell cell type gene signatures (gsQTLs) in TCGA.
6. Shahamatdar et al. *Cell Reports* 2020. Systematic identification of germline genetic polymorphisms associated with immune infiltration in TCGA.
7. Ostendorf et al. *Nat Medicine* 2020. Identification of *APOE2* and *APOE4* germline variants associated with melanoma progression and ICB response in mice.
8. Zhang et al. *Front Genet* 2019. Identification of breast-cancer associated variant modulating *CTSW* expression
9. Sayaman et al. *Immunity* 2020. Systematic identification of germline variants associated with 33 immune traits including leukocyte subsets, adaptive receptor, immune expression signatures.
10. Yoshida et al. *Eur J Cancer* 2021. Identification of 2 *PD-L1* variants associated with survival outcomes in advanced non-small-cell lung cancer patients.
11. Kula et al. *Exp Mol Pathol* 2020. Review of 10 *PD-L1* genetics variants.
12. Salmaninejad et al. *Immunogenetics* 2018. Review of 5 frequently studied *PD-1* genetic variants.
13. Sasaki et al. *Mol Clin Oncol* 2014. Characterization of *PD-1* promoter variant and association with survival in non-small cell lung cancer.
14. Tang et al. *Int J Clin Exp Med* 2015. Characterization of 3 *PD-1* variants and association with cancer risk.

For Sayaman et al., 598 significant associations were identified, 520 of which were within the MHC II region. To identify independent Sayaman et al SNPs, we performed linkage disequilibrium based clumping with the same parameters used for our analysis. After clumping, 55 independent Sayaman et al snps remained.

### GREGOR (RRID: SCR_009165)

GREGOR (RRID: SCR_009165) was used to analyze SNP enrichment at epigenetic features. We obtained 479 bed files for 11 histone experiments and 52 cell types from ENCODE (RRID:SCR_015482) (downloaded on May 3, 2020). Only “stable peaks” and “replicated peaks” files were kept for analysis. If more than 1 bed file for a cell type and histone mark were available, the files were combined.

In addition, 323 bed files for 12 transcription factor binding experiments and 12 cell types were downloaded from ENCODE (RRID:SCR_015482) on August 4, 2020. Only “optimal IDR thresholded peaks” and “conservative IDR thresholded peaks” files were kept. If more than 1 bed file for a cell type and transcription factor were available, the files were combined.

GREGOR (RRID: SCR_009165) was run with EUR Reference files made from the 1000 Genomes Project data with an LD window size of 1MB and LD R^2^ > 0.7. Enrichment ratios were calculated by taking the difference between observed and expected number of SNPs and dividing by the expected number of SNPs. Any files with Audit errors were excluded.

### Variant Annotation

Variants were annotated with VEP (Variant Effect Predictor)^133^ with default parameters and the GRCh37 reference genome. Coding variants were mapped to protein sequences using the Uniprot GFF file.

### Cell-Type eQTL Analysis

We followed the GTEx approach for cell type interaction eQTL discovery^31^. We ran a linear regression model with an interaction term accounting for interactions between genotype and cell type enrichment from xCell^134^ **Eqn 2**:

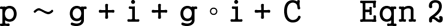

where p is the IP component vector, g is the genotype vector, i is the inverse normal transformed by tissue type xCell enrichment score^134^, and the interaction term g ◦ i corresponds to pointwise multiplication of genotypes and cell type enrichment scores. The same covariates, denoted by C, were used as in the regular immune microenvironment GWAS analysis. Benjamini-Hochberg FDR was calculated for the beta coefficient of the interaction term and variants with FDR < 0.1 were identified as significant.

DICE expression quantitative trait loci (eQTLs) were obtained at https://dice-database.org/. Methods associated with DICE eQTL discovery are published in Schmiedel et al^135^.

### UK Biobank

UK Biobank subjects were subsetted into separate ethnic-racial groups following continental ancestry prior to analysis. The sub-setting was performed to generate homogenous groups and reduce potential admixture bias in the genetic analyses. To identify the European-ancestry samples, we started with directly called genotype data and identified a set of overlapping SNPs with 1000 Genomes Project and AWS (RRID:SCR_008801) (1KG) population and then merged them together. Next, we pruned the SNP set so remaining SNPs were in linkage equilibrium using PLINK (PLINK, RRID: SCR_001757)^30^. flashpca was used to calculate principal components for 1KG SNPs^136^. The UK Biobank samples were projected onto 1KG space using flashpca. To identify subjects of European ancestry, we utilized Aberrant to generate clusters with a broad set of lambda values (clustering thresholds) and checked that the cluster included all 1KG subjects of European ancestry and maximized the total number of UK Biobank subjects (lambda=8.2)^124^. Finally, we compared the self-reported race/ethnicity of subjects within this cluster and removed samples that were discordant. We identified 454,487 subjects of European ancestry. To identify the unrelated samples from the finalized European list, we used the relatedness file provided by UK Biobank and a custom script was used to select unrelated samples while maximizing sample counts. The final European unrelated set included 382,841 subjects.

Variant dosages extracted from imputed UK Biobank BGEN files were used for PheWAS analysis with PLATO v2.0.0^137^. ICD10 diagnosis codes associated with neoplasms and immune disorders were collapsed according to level-1 groupings used by UK Biobank resulting in a total of 24 groups. For example, C00-C14 is one of the groups containing ICD10 codes associated with malignant neoplasm of lip, oral cavity, and pharynx. Individuals with diagnosis code in a group were coded as 1, with the remaining individuals coded as 0. Logistic regression was conducted with UK Biobank binary files containing HLA-immune variants, logistic phenotype file, and age, sex, and principal components 1-10 as covariates. Pvalues were Benjamini-Hochberg FDR adjusted.

### Survival Analysis

Kaplan-Meier analysis of immune microenvironment associations were conducted with overall and progression- free survival retrieved from Liu et al.^141^ by cancer type using the lifelines package. As recommended by Liu *et al,* TCGA cancer types, TGCT and PCPG, were excluded as survival data did not meet quality standards. TCGA individuals were divided into 3 groups based on genotype calls: minor allele homozygotes, heterozygotes and major allele homozygotes. Significance was determined using the logrank test between minor allele and major allele homozygotes. Only SNPs with at least 1% minor allele frequency in each cancer type and more than 1 minor allele homozygous individual were considered for analysis. Logrank p values were corrected using the Benjamini-Hochberg method from the statsmodel package and only variants with FDR < 0.1 were considered.

### Immune Checkpoint Blockade (ICB) Cohort Genotypes

Raw fastq files were obtained for the following immune checkpoint trials: Hugo et al. 2016 (SRA accession: SRP090294, SRP067938; Cancer: melanoma)^51^, Van Allen et al. 2015 (SRA accession: SRP011540, Cancer: melanoma)^52^, Miao et al. 2018 (SRA accession: SRP128156, Cancer: clear cell renal carcinoma)^138^, Riaz et al. 2017 (SRA accession: SRP095809, SRP094781; Cancer: melanoma)^54^, Rizvi et al. 2015 (SRA accession: SRP064805, Cancer: non-small cell lung cancer)^53^, Snyder et al. 2014 (SRA accession: SRP072934, Cancer: melanoma)^55^. Reads were aligned to UCSC hg19 coordinates using BWA (RRID:SCR_010910) v0.7.17-r1188^139^. Reads were sorted by SAMTOOLS (RRID:SCR_002105) v0.1.19^140, 141^, marked for duplicates with Picard Tools (RRID:SCR_006525) v2.12.3 and recalibrated with GATK (RRID:SCR_001876) v3.8-1-0^142–144^. Germline variants were called from sorted BAM files using DeepVariant v0.10.0-gpu^145, 146^. The final immunotherapy cohort consisted of 68 clear cell renal carcinoma, 276 melanoma and 34 non-small cell lung cancer patients.

To evaluate the quality of SNP imputation from whole exome data, we took advantage of the TCGA having both. Of the 1,322,586 variants available from DeepVariant analysis of immunotherapy cohort, 225,000 were available in TCGA imputed data. We extracted these 225,000 variants from TCGA and input into the Michigan Imputation Server (reference panel: HRC, phasing: Eagle). We compared genotypes from whole-exome calls vs. original Affymetrix-based TIME-SNP calls. Variants with >5% mismatches in genotype calls, minor allele frequency < 5% in any cohort or imputation accuracy (R^2^ < 0.3) were excluded. Only variants with at least 5% frequency in all 4 melanoma cohorts used for discovery analysis were considered for ICB analysis, leaving 525 SNPs.

Population stratification analysis was conducted by taking overlapping variants between TCGA and ICB cohorts. Variants with MAF differences > 0.1% were excluded resulting in 3612 frequency-concordant variants. PLINK IBD analysis was conducted and top 10 principal components were included in association analysis.

### Immune Checkpoint Blockade (ICB) Response Analysis

Subject phenotypes were downloaded from supplementary information of ICB trial publications. Four melanoma cohorts were used as the discovery cohort for ICB-associated variants, while Miao et al renal cell carcinoma and Rizvi et al non-small cell lung cancer cohorts were used for validation. Response phenotypes were determined from iRECIST criteria^147^. Patients were categorized as responders if they had iRECIST criteria: CR (complete response), PR (partial response) and SD (stable disease). Genome-wide association studies (GWASs) were conducted for ICB responders within each ICB-cohort using PLINK (PLINK, RRID:SCR_001757). Age, sex and the top 10 principal components were included in the logistic analysis as covariates. We then used METAL^58^ with a sample size weighting scheme to perform a pan-study melanoma meta-analysis for ICB response.

Eleven variants (FDR < 0.5) were selected for construction of a germline burden score. Variant genotypes were aligned such that the allele associated with higher odds of responder status were used for construction of the burden score, which was a simple count of the number of ICB-associated variants for each individual.

### Immune Checkpoint Blockade (ICB) Response RNA-seq

FASTQ/BAM files were downloaded for 33 RCC and 120 melanoma patients. BAM files were converted to FASTQ using bam2fq^141^. Unpaired reads were removed using fastq pair^148^. Paired reads were aligned with STAR (RRID:SCR_004463) v2.4.1d^149^ to GRCh37 reference alignment. RSEM^150^ was used for transcript quantification. TPM values were log2 transformed for analyses. Differential gene expression analysis between responders and non-responders from cohorts Riaz et al. 2017, Hugo et al. 2016, Miao et al. 2018, and Van Allen et al. 2015 was performed using the DESeq2^143^ package in R. Cohort was included as a covariate when calculating top differentially expressed genes.

### Mouse experiments

Wild-type C57BL/6 (RRID:IMSR_JAX:000664) were purchased from The Jackson Laboratory. All the animal studies were approved by the Institutional Animal Care and Use Committee (IACUC) of university of California, San Diego, with protocol ASP #S15195. Mice at Moores Cancer Center, UCSD are housed in micro- isolator and individually ventilated cages supplied with acidified water and fed 5053 Irradiated Picolab Rodent Diet 20 lab diet. Temperature for laboratory mice in our facility is mandated to be between 65–75 ° F (∼18–23 °C) with 40–60% humidity. All animal manipulation activities are conducted in laminar flow hoods. All personnel are required to wear scrubs and/or lab coat, mask, hair net, dedicated shoes, and disposable gloves upon entering the animal rooms. 2x10^5^ MC38 (RRID:CVCL_B288) cells were transplanted into the flank of 8-10 female C57Bl/6 (RRID:IMSR_JAX:000664) mice, aged 7-8 weeks. Where indicated, when tumors reached 100 mm^3^, mice were randomized and treated with anti-PD-1 (10mg/kg i.p., Bio X Cell Cat# BE0146, RRID: AB10949053, clone RMP1-14), CTSS inhibitor (5mg/kg, i.p., APEx Bio) or isotype control antibody. Treatments were given 3 times a week. Where indicated (or when control-treated mice succumbed to tumor burdens, as determined by the ASP guidelines), mice were euthanized, and tumors were taken for flow cytometric analysis. MC38 cells were not screened using STR profiled on site.

### Reagents

PD-1 antibody (clone RMP1-14, Bio X Cell Cat# BE0146, RRID:AB_10949053) and isotype antibody (catalog #BE0091) were purchased from Bio X Cell.

### RT-PCR

RNA from MC38 (RRID:CVCL_B288) tumors was extracted using the RNeasy Mini Kit (Qiagen catalog #74104). 500ng of RNA per reaction was used to prepare cDNA with the SuperScript™ VILO™ cDNA Synthesis Kit (ThermoFisher Scientific) following manufacturer’s instructions. The cDNA was used to set up the RT-PCR reaction with 4 technical replicates per tumor with the Fast SYBR™ Green Master Mix (ThermoFisher Scientific) according to manufacturer’s instructions. PCR quantification was conducted using the 2^-ΔΔCT^ method and normalized to the housekeeping gene β-actin. Primers used for *CTSS* expression quantification is detailed in **Table S3**.

RNA-seq and CIBERSORTx infiltration estimates for M4 melanoma mouse model were obtained from GEO accession (GSE144946). Responders were mice whose size at harvest was smaller than the last dose of anti-CTLA-4. RNA-seq counts were converted to TPM and log2 normalized.

