## Extended Figures/Tables for "Germline modifiers of the tumor immune microenvironment implicate drivers of cancer risk and immunotherapy response"

| **TCGA Cancer type abbreviation** | **Cancer** |
|---|---|
| LUSC | Lung squamous cell carcinoma |
| LUAD | Lung adenocarcinoma |
| KIRC | Kidney renal cell carcinoma |
| KIRP | Kidney renal papillary cell carcinoma |
| COAD | Colon adenocarcinoma |
| BRCA | Breast invasive carcinoma |
| READ | Rectum adenocarcinoma |
| UCEC | Uterine corpus endometrial carcinoma |
| LIHC | Liver hepatocellular carcinoma |
| THCA | Thyroid carcinoma |
| BLCA | Bladder urothelial carcinoma |
| STAD | Stomach adenocarcinoma |
| PRAD | Prostate adenocarcinoma |
| HNSC | Head and Neck squamous cell carcinoma |
| CESC | Cervical squamous cell carcinoma and endocervical adenocarcinoma |
| SARC | Sarcoma |
| SKCM | Skin cutaneous melanoma |
| PAAD | Pancreatic adenocarcinoma |
| ESCA | Esophageal carcinoma |
| KICH | Kidney chromophobe |
| PCPG | Pheochromocytoma and Paraganglioma |
| CHOL | Cholangiocarcinoma |

**Table S1: TCGA Cancer Types**

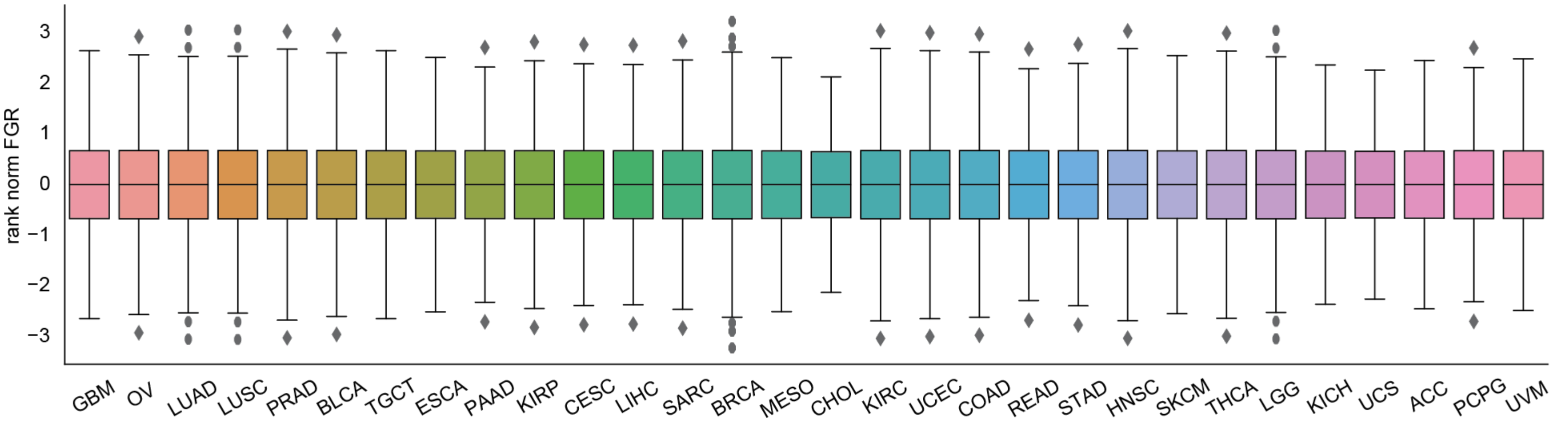


**Figure S1: Characterization of Tumor Immune Microenvironment, Related to Figure 1.** Boxplot of FGR expression values after inverse-rank normalization by cancer type.

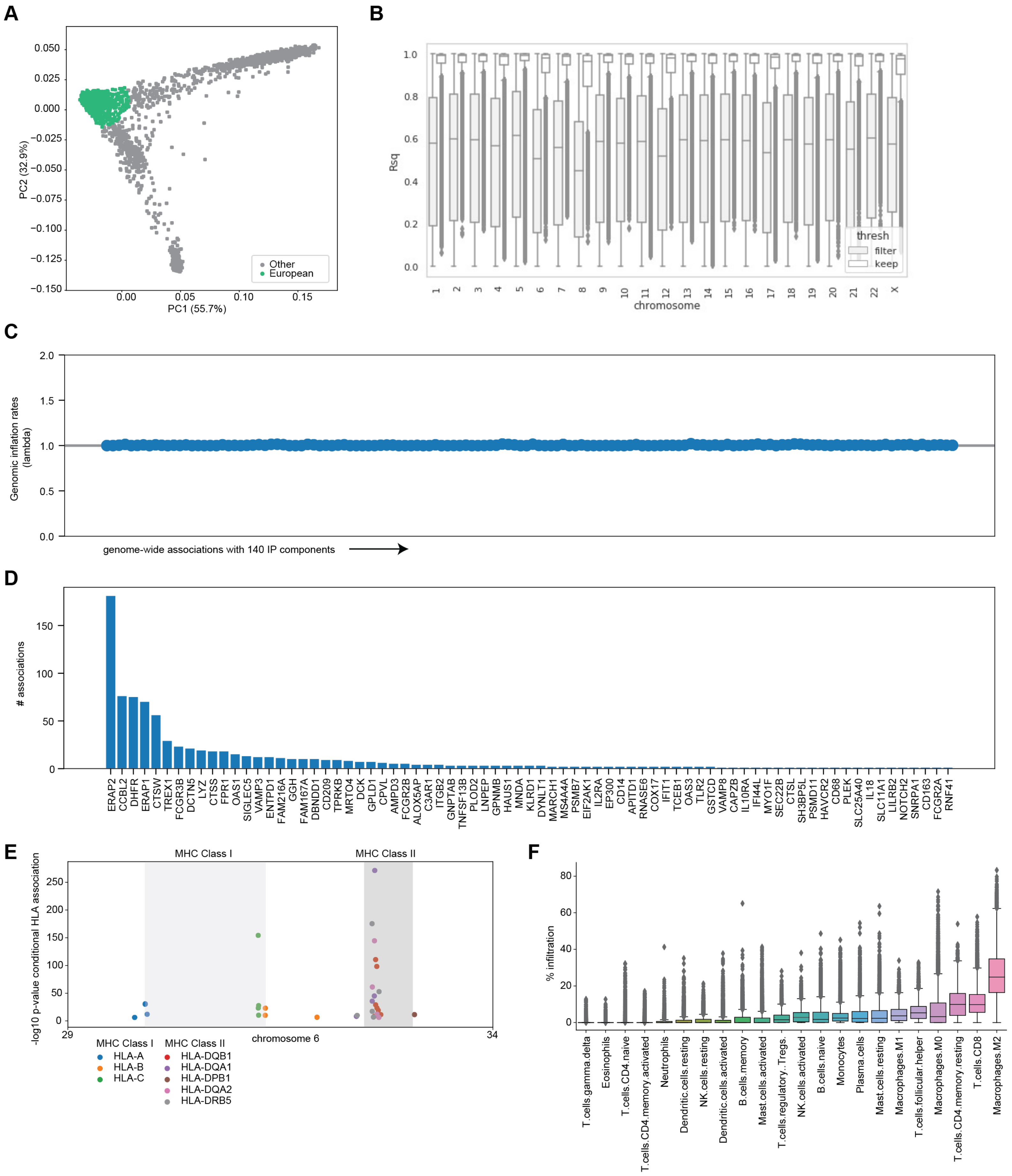

A
PC1 (55.7%)
PC2 (32.9%)
Other
European
B
Rsq
chromosome
thresh
filter
keep
C
Genomic inflation rates (lambda)
genome-wide associations with 140 IP components
D
### associations
ERAP2 CCBL2 DHFR ERAP1 CTSW TREX1 FCGR3B DCTN5 LYZ CTSS FPR1 OAS1 SIGLEC5 VAMP3 ENTPD1 FAM216A GGH FAM167A DBNDD1 CD209 TPRKB MRTO4 DCK GPLD1 CPVL AMPD3 FCGR2B ALOX5AP C3AR1 ITGB2 GNPTAB TNFSF13B PLOD2 LNPEP GPNMB HAUS1 MNDA KLRD1 DYNLT1 MARCH1 MS4A4A PSMB7 EIF2AK1 IL2RA EP300 CD14 APITD1 RNASE6 COX17 IFIT1 TCEB1 OAS3 TLR2 GSTCD VAMP8 CAPZB IL10RA IFI44L MYO1F SEC22B CTSL SH3BP5L PSMD11 HAVCR2 CD68 PLEK SLC25A40 IL18 SLC11A1 LILRB2 NOTCH2 SNRPA1 CD163 FCGR2A RNF41
E
MHC Class I
MHC Class II
-log10 p-value conditional HLA association
chromosome 6
HLA-A
HLA-B
HLA-C
HLA-DQB1
HLA-DQA1
HLA-DPB1
HLA-DQA2
HLA-DRB5
F
% infiltration
T.cells.gamma.delta Eosinophils T.cells.CD4.naive T.cells.CD4.memory.activated Neutrophils Dendritic.cells.resting NK.cells.resting Dendritic.cells.activated B.cells.memory Mast.cells.activated T.cells.regulatory..Tregs. NK.cells.activated B.cells.naive Monocytes Plasma.cells Mast.cells.resting Macrophages.M1 T.cells.follicular.helper Macrophages.M0 T.cells.CD4.memory.resting T.cells.CD8 Macrophages.M2

**Figure S2: Detecting putative germline modifiers of the tumor immune microenvironment , Related to Figure 2. (A).** PCA analysis of TCGA genotypes; green indicates European discovery cohort. **(B)** Boxplot of Rsq distribution of imputed genotypes across chromosomes. **(C)** Plot of inflation factors (lambda) for associations using PancanAtlas data. **(D)** Barplot of number of associations for each immune phenotype (IP) components. **(E)** Scatterplot of HLA associations based on chromosome 6 genomic location. **(F)** Boxplot of CIBERSORTx infiltration across TCGA individuals.

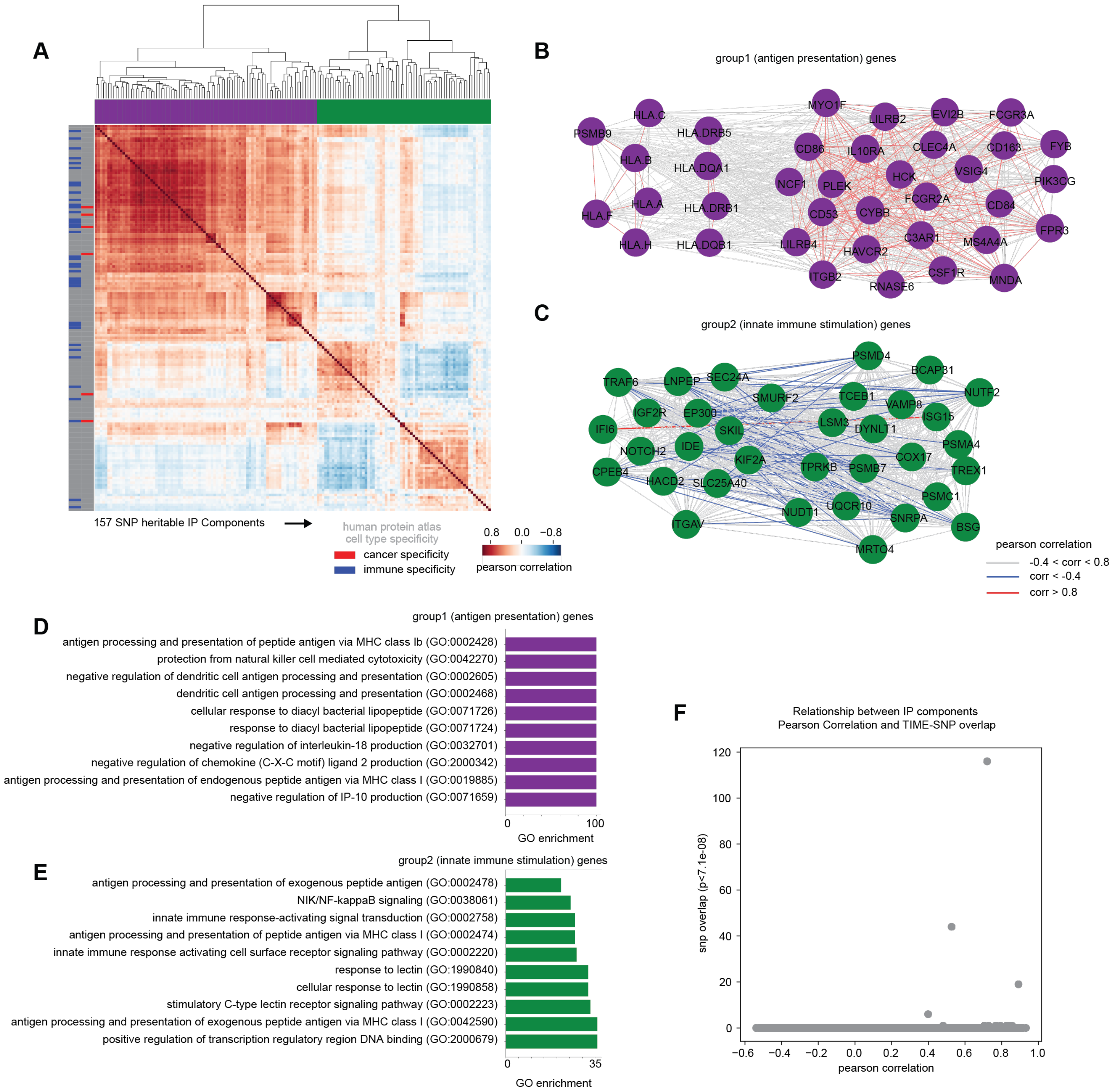


**Figure S3: Detecting putative germline modifiers of the tumor immune microenvironment, Related to Figure 2. (A)** Clustermap depicting 157 SNP-heritable IP components from two-state GCTA analysis and their pairwise correlation across 30 tumors in the TCGA. **(B)** Network plot of group 1 (antigen presentation) genes from pearson correlation Clustermap analysis. **(C)** Network plot of group 2 (innate immune stimulation) genes from pearson correlation Clustermap analysis **(D)** Top 10 GO enrichment terms and enrichment values from group 1 (antigen presentation) genes. **(E)** Top 10 GO enrichment categories and enrichment values from group 2 (innate immune stimulation) genes. **(F)** Scatterplot of pearson correlation of IP components and number of overlapping significant variants.

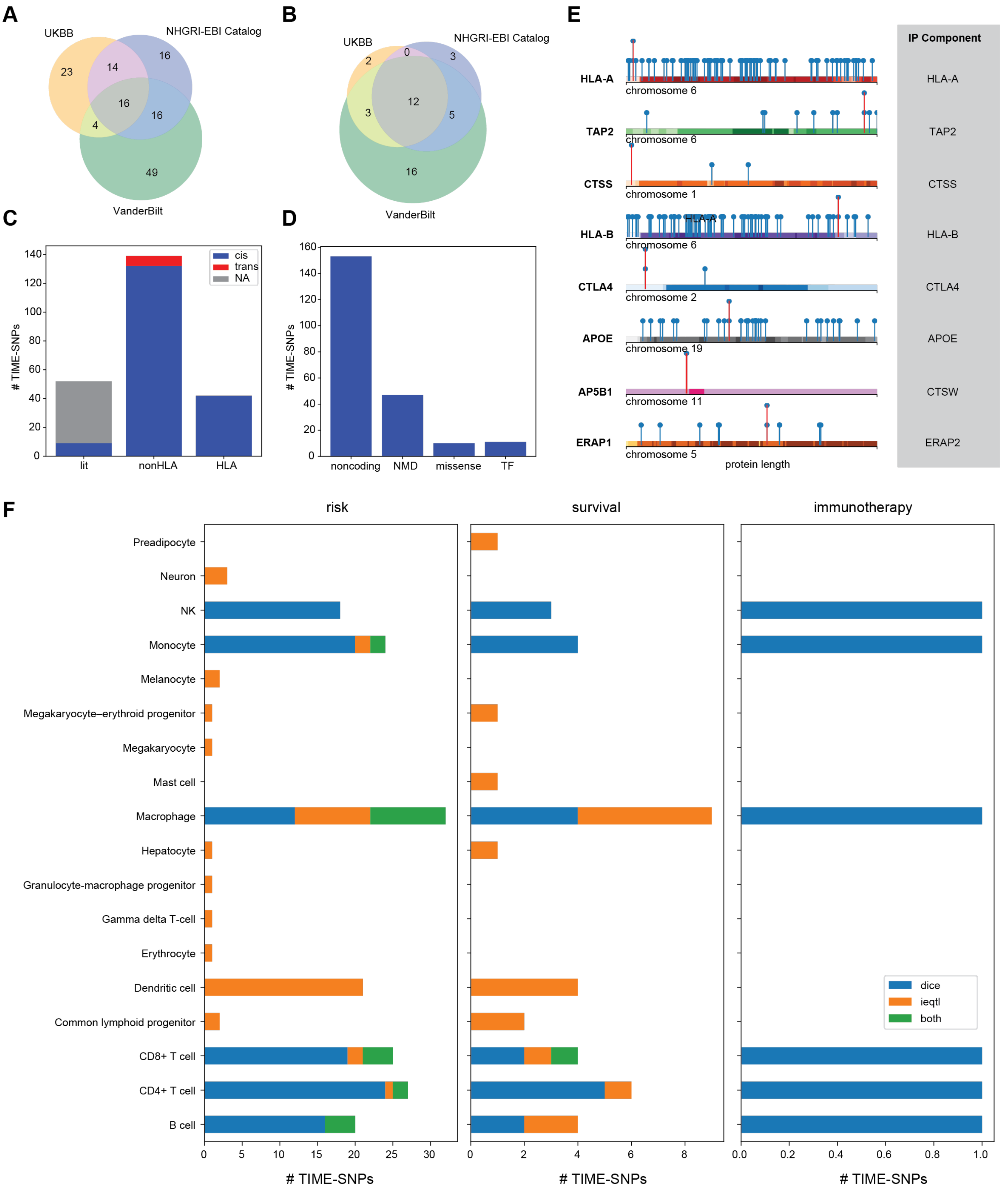
A
UKBB
NHGRI-EBI Catalog
VanderBilt
23
14
16
16
16
4
49
B
UKBB
NHGRI-EBI Catalog
VanderBilt
2
0
3
12
3
5
16
C
### TIME-SNPs
cis
trans
NA
lit
nonHLA
HLA
D
### TIME-SNPs
noncoding
NMD
missense
TF
E
IP Component
HLA-A
TAP2
CTSS
HLA-B
CTLA4
APOE
AP5B1
ERAP1
chromosome 6
chromosome 6
chromosome 1
chromosome 6
chromosome 2
chromosome 19
chromosome 11
chromosome 5
protein length
HLA-A
TAP2
CTSS
HLA-B
CTLA4
APOE
CTSW
ERAP2
F
risk
survival
immunotherapy
Preadipocyte
Neuron
NK
Monocyte
Melanocyte
Megakaryocyte–erythroid progenitor
Megakaryocyte
Mast cell
Macrophage
Hepatocyte
Granulocyte-macrophage progenitor
Gamma delta T-cell
Erythrocyte
Dendritic cell
Common lymphoid progenitor
CD8+ T cell
CD4+ T cell
B cell
dice
ieqtl
both
### TIME-SNPs
### TIME-SNPs
### TIME-SNPs

**Figure S4: Identification and characterization TIME-SNPs related to cancer outcomes, Related to Figure 3. (A)** Venn Diagram of overlap between TIME-SNPs implicated by UK Biobank PheWAS, Vanderbilt PheWAS catalog and NHGRI-EBI GWAS Catalog (LDtrait). **(B)** Venn Diagram of overlap between IP Components implicated by TIME-SNPs by UK Biobank PheWAS, Vanderbilt PheWAS catalog and NHGRI-EBI GWAS Catalog. **(C)** Barplot quantifying the number of cancer relevant TIME-SNPs which are *trans* (>1 MB from the TSS of the associated IP component) and *cis*. **(D)**. Barplot describing the effects of cancer relevant TIME-SNPs. **(E)** Location of missense TIME-SNPs (red) and known genetic variation (blue) in the coding sequence of affected IP components. The IP component with which the SNP was associated is shown on the right side. **(F)** Boxplot of number of cancer risk, survival and immunotherapy response TIME-SNPs which are cell-type eQTLs.

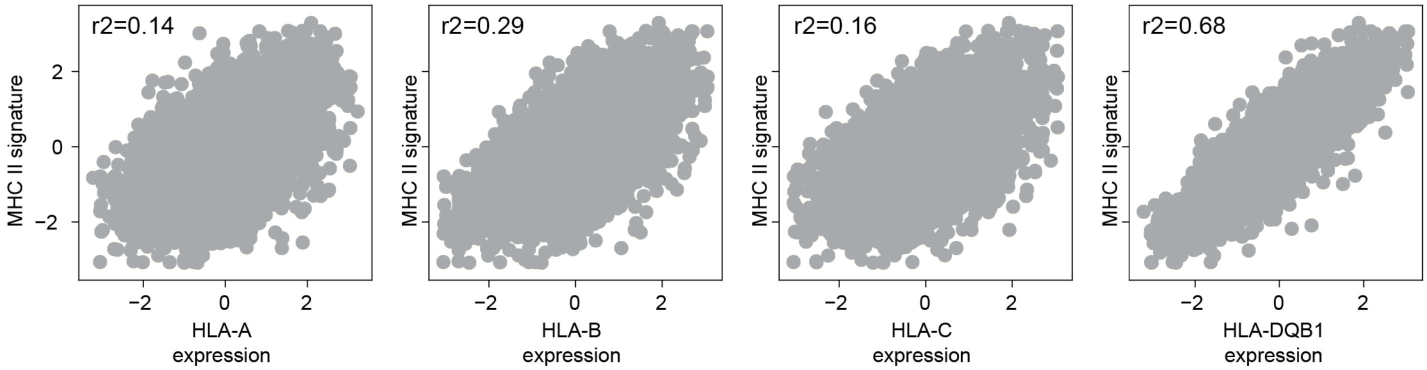


**Figure S5: TIME-SNPs underlying antigen presentation stratify melanoma and prostate cancer risk, Related to Figure 4.** Correlation between Sayaman et al. MHC Class II signature and HLA-A, HLA-B, HLA-C and HLA-DQB1 expression.

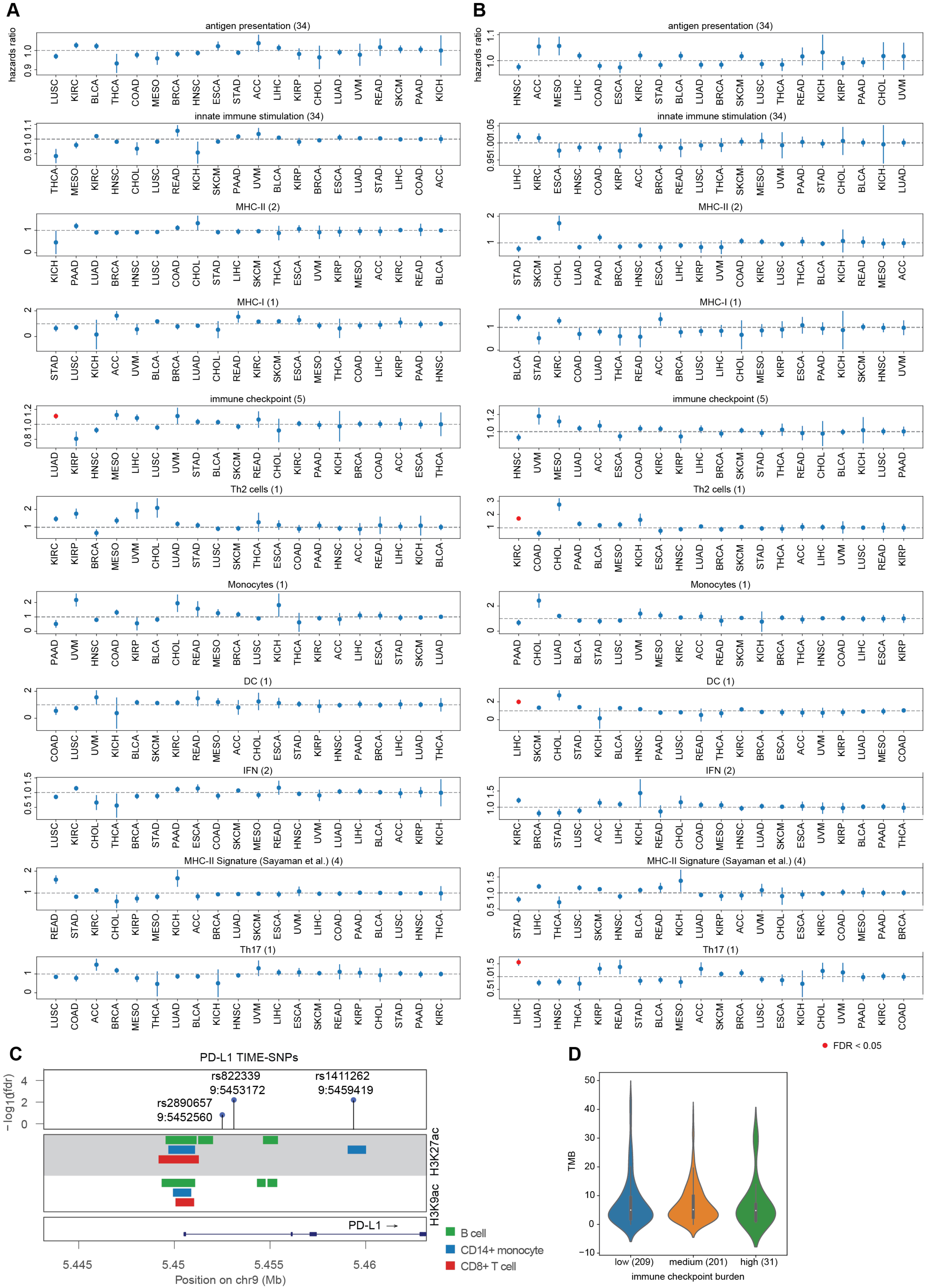

A
B
hazards ratio
antigen presentation (34)
innate immune stimulation (34)
MHC-II (2)
MHC-I (1)
immune checkpoint (5)
Th2 cells (1)
Monocytes (1)
DC (1)
IFN (2)
MHC-II Signature (Sayaman et al.) (4)
Th17 (1)
FDR < 0.05
C
PD-L1 TIME-SNPs
−log10(fdr)
rs822339
9:5453172
rs1411262
9:5459419
rs2890657
9:5452560
H3K27ac
H3K9ac
PD-L1 →
5.445
5.45
5.455
5.46
Position on chr9 (Mb)
B cell
CD14+ monocyte
CD8+ T cell
D
TMB
low (209)
medium (201)
high (31)
immune checkpoint burden

**Figure S6: Variants underlying immune Evasion are associated with cancer survival, Related to Figure 5. (A)** Cox Proportional-Hazards results from association with burden score, if more than 1 variant association for IP category was detected, and overall survival for each TCGA cancer type. **(B)**. Cox Proportional-Hazards results from association with burden score, if more than 1 variant association for IP category was detected, and progression-free survival for each TCGA cancer type. **(C)** Location of PD-L1 variants implicated through literature review in relation to cell-type specific H3K27ac and H3K9ac marks. **(D)** Violin plot of immune checkpoint burden score and tumor mutational burden.

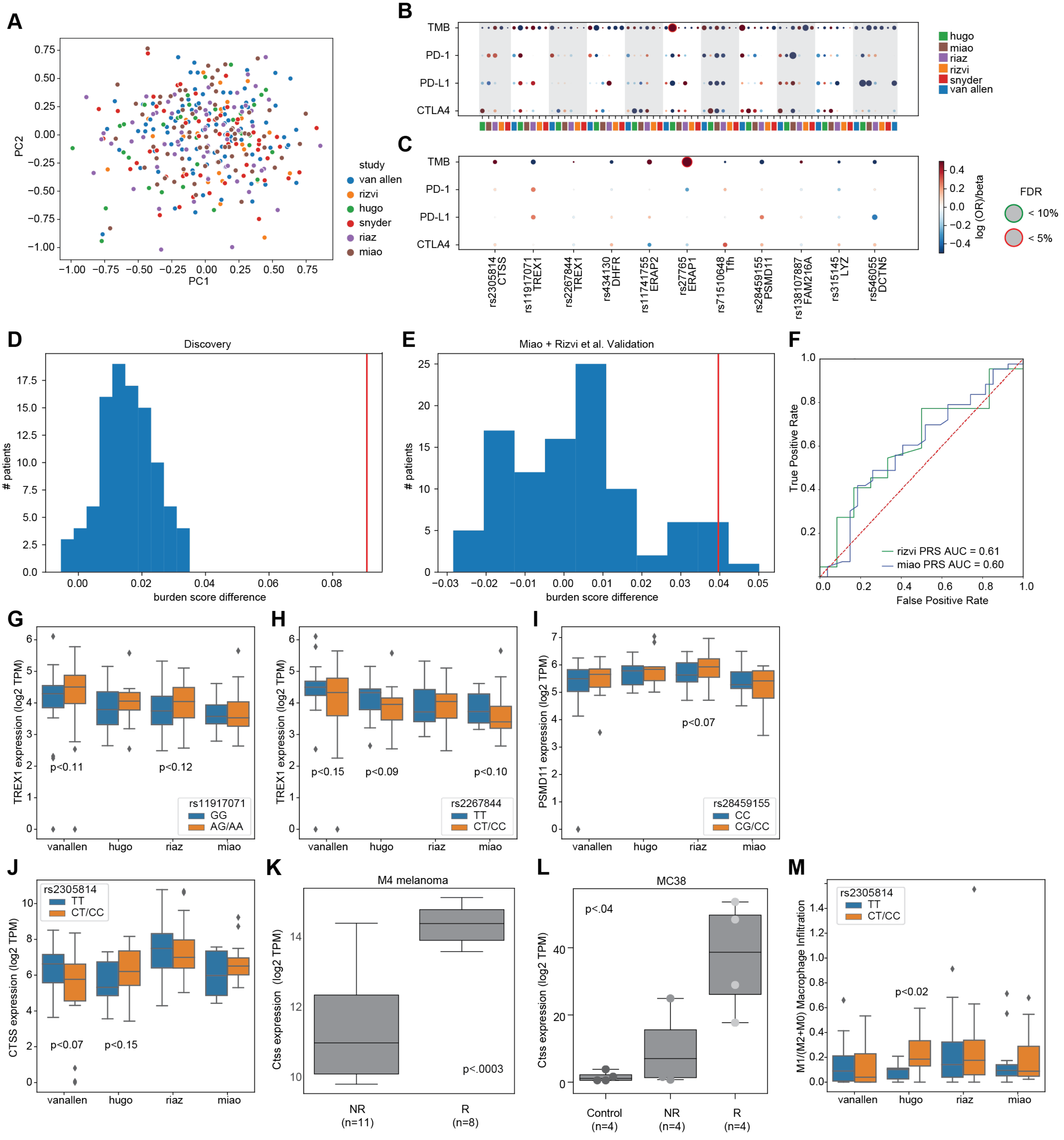

A
B
C
D
Discovery
E
Miao + Rizvi et al. Validation
F
G
H
I
J
K
M4 melanoma
L
MC38
M
PC1
PC2
study
van allen
rizvi
hugo
snyder
riaz
miao
TMB
PD-1
PD-L1
CTLA4
log (OR)/beta
FDR
< 10%
< 5%
rs2305814 CTSS
rs11917071 TREX1
rs2267844 TREX1
rs434130 DHFR
rs11741755 ERAP2
rs27765 ERAP1
rs71510648 Tfh
rs28459155 PSMD11
rs138107887 FAM216A
rs315145 LYZ
rs546055 DCTN5
### patients
burden score difference
True Positive Rate
False Positive Rate
rizvi PRS AUC = 0.61
miao PRS AUC = 0.60
TREX1 expression (log2 TPM)
p<0.11
p<0.12
rs11917071
GG
AG/AA
p<0.15
p<0.09
p<0.10
rs2267844
TT
CT/CC
PSMD11 expression (log2 TPM)
p<0.07
rs28459155
CC
CG/CC
vanallen
hugo
riaz
miao
CTSS expression (log2 TPM)
rs2305814
TT
CT/CC
p<0.07
p<0.15
Ctss expression (log2 TPM)
p<.0003
NR (n=11)
R (n=8)
p<.04
Control (n=4)
NR (n=4)
R (n=4)
M1/(M2+M0) Macrophage Infiltration
p<0.02

**Figure S7: TIME-SNPs implicate targets for modulating immune responses, Related to Figure 6.**
(**A)** PCA plot of 11 TIME-SNPs associated with immunotherapy response (FDR<0.5) **(B)** Grid plot of beta values of variant association with TMB, *CTLA-4, PD-1*, and *PD-L1* by each cohort. **(C)** Grid plot of beta values of variant association with TMB, *CTLA-4, PD-1*, and *PD-L1* controlling for cohort **(D)** Histogram of difference between average burden score for 100 random bootstrapping trials compared to actual difference based on 11 TIME-SNPs (red) in discovery cohort. **(E)** Histogram of difference between average burden score for 100 random bootstrapping trials compared to actual difference based on 11 TIME-SNPs (red) in Miao and Rizvi et al validation cohort. **(F)** ROC-AUC Curve Analysis for PRSice PRS scores trained on the discovery cohort and tested on Miao et al and Rizvi et al. **(G)** Boxplot of rs11917071 association with *TREX1* expression across ICB cohorts. **(H)** Boxplot of rs2267844 association with *TREX1* expression across ICB cohorts. **(I)** Boxplot of rs28459155 association with *PSMD11* expression across ICB cohorts. **(J)** Boxplot of rs2305814 association with *CTSS* expression across ICB cohorts. **(K)** *Ctss* expression in non-responder and responder of M4 melanoma mouse model. **(L)** *Ctss* expression in non-responder and responder of MC38 mouse model. **(M)** Boxplot of rs2305814 association with M1/M2+M0 macrophage infiltration across ICB cohorts.

| CTSS | |
|---|---|
| Forward Primer | CCATTGGGATCTCTGGAAGAAAA |
| Reverse Primer | TCATGCCCACTTGGTAGGTAT |

**Table S2: qPCR primers for *Ctss* quantification in MC38 mouse model**
